# Comprehensive LC-MS/MS Data Acquisition in Metabolomics via Maximum Bipartite Matching

**DOI:** 10.1101/2025.02.22.639652

**Authors:** Ross McBride, Stefan Weidt, Joe Wandy, Vinny Davies, Rónán Daly, Kevin Bryson

## Abstract

**Background:** In untargeted metabolomics studies, liquid chromatography tandem mass spectrometry (LC-MS/MS) is a powerful analytical platform. The fragmentation spectra produced can be used as “molecular fingerprints” to identify unknown metabolites. However, the high number of analytes that may be co-eluting limits the number of fragmentation spectra that can be collected and potentially identified, presenting a serious bottleneck for many studies. There is a need for new fragmentation strategies which are comprehensive, interpretable and robust, meaning they produce high-quality fragmentation spectra for as many analytes as possible while operating within the constraints of notoriously noisy mass spectrometry data.

**Results:** We present a data acquisition workflow which uses a bipartite graph to represent the relationship between opportunities for fragmentation and desired fragmentation targets. This method allows a schedule for data acquisition to be optimally allocated by a standard algorithm. We augment this existing technique by allowing it to solve for multiple samples collectively, allowing it to optimise target intensity (and hence spectral quality) via the use of a weighted matching and by assigning leftover scans redundantly to improve robustness. We also show how this workflow can be used flexibly to generate inclusion windows for Data-Dependent Acquisition (DDA) methods. Our experiments show that several thousand peaks identified in a realistic biological sample can be targeted using only two LC-MS/MS runs. We also further investigate the trade-off between offline workflows and DDA methods by exposing our target list of peaks to realistic variation across samples. We find in those circumstances that our new method has performance (measured by number of peaks targeted comparable to state-of-the-art DDA methods). However, this competitive performance is only possible with our additions to the base maximum matching technique, which provide extra resistance against inter-sample variations.

**Conclusions:** We have proposed a workflow for LC-MS/MS data acquisition which can be used flexibly for entirely pre-scheduled acquisition or which may generate inclusion windows for online DDA methods. Our results show that the maximum matching workflow with our improvements is state-of-the-art where pre-scheduling is concerned, and in future this foundation may be developed to build more powerful DDA methods which can action the promise of truly comprehensive data acquisition.

## 1 Introduction

Untargeted metabolomics experiments attempt to profile the metabolome of a biological sample as completely as possible. A key technology in this analysis is liquid chromatography tandem mass spectrometry (LC-MS/MS) [23, 21]. In mass spectrometry (MS) a chemical sample is ionised, ions are separated by their mass-to-charge ratio (m/z) and a detector measures their intensity, from which their relative abundance can be inferred. With the use of electrospray ionisation this technique can be coupled to liquid chromatography (LC), which, prior to ionisation, further separates analytes (including those of equal m/z) by retention time (RT). Measurement of these (RT, m/z, intensity) triplets at a given RT is called a “survey” or “MS1” scan. Individual analytes produce traces along RT (and therefore across multiple MS1 scans) known as “chromatographic peaks”. With the addition of tandem mass spectrometry (MS/MS), we can also isolate ions of a specific m/z range, fragment them and measure (m/z, intensity) pairs. The “fragmentation spectra” produced by these “fragmentation” or “MS2” scans offer information on how “precursor” ions captured by the isolation window broke apart. They therefore act as a kind of “molecular fingerprint”, allowing chemical structure to be inferred. An untargeted metabolomics experiment therefore desires high-quality fragmentation spectra for as many unique analytes as possible. However, this is a difficult problem in practice and presents a major bottleneck for many metabolomics studies [15, 20].

The balance of MS1 and MS2 scans, and the target m/z window for individual MS2 scans, are controlled by a fragmentation strategy. A good choice of fragmentation strategy can therefore aid in reducing this bottleneck on metabolomics studies. A good fragmentation strategy should satisfy (at least) the following criteria:

- Comprehensiveness: fragmentation spectra should be obtained for as many unique analytes as possible.
- Interpretability: fragmentation spectra should be of sufficient quality that any relevant chemical information can be inferred from them.
- Robustness: the strategy should still function under conditions of noisy data.

The most well-established fragmentation strategy is the Data-Dependent Acquisition (DDA) method, **TopN** [5, 31]. TopN is a simple heuristic where an MS1 “survey scan” is followed by up to *N* MS2 scans, and MS2 scans are targeted at *m/z* windows centred on the *N* highest intensity peaks in the preceding MS1 scan. Potential targets must exceed a fixed intensity threshold and less than *N* MS2 scans will be scheduled if there are insufficient targets. Although the most intense ions are often most relevant to sample characterisation, profiling is not very comprehensive as TopN repeatedly fragments the same high-intensity ions. To mitigate this behaviour TopN can be augmented by exclusion/inclusion windows which specify an interval in (*rt, m/z*) space to avoid/prioritise. These can be loaded into an MS instrument at the start of a run, but a Dynamic Exclusion Window (DEW) can also be created on every fragmentation event to prohibit refragmentation within a fixed (*rt, m/z*) window.

As it targets individual ions according to naive heuristics, TopN is typically not very comprehensive even with standard modifications like DEWs. This shortcoming has generated research interest in Data Independent Acquisition (DIA). DIA methods use a scheme of large isolation windows with no real-time feedback. Each window captures many ions, so DIA methods are often entirely comprehensive, but this then requires combined spectra to be deconvolved or otherwise interpreted. Deconvolution is difficult [9, 30] and is an area of ongoing research [26, 24]. But strategies which target individual ions, like TopN, can usually attribute all fragments to a single parent ion, offering easy interpretability. Therefore an alternative, parallel research direction is to improve the comprehensiveness of such methods.

**TopN Exclusion** [1, 16] remembers DEWs between runs, using them as regular exclusion windows in future runs. This forces TopN to progressively explore signals in different regions of (*rt, m/z*) space, increasing its comprehensiveness. **WeightedDEW** [4] replaces the binary exclusion of the standard DEW with a DEW split into a binary exclusion and a linear weighting of intensity based on RT. **SmartRoI** [4] does not use regular DEWs, instead building Regions-of-Interest (RoIs) around chromatographic traces in real-time and performing binary exclusion on an RoI if its intensity has neither risen nor fallen by a specified factor since its last fragmentation. Both were shown to increase sample coverage compared to TopN. TopNEXt [16] is a framework for DDA methods which generalises these methods into a system of scoring functions and enables more advanced manipulation of RoIs. The best-performing method implemented via TopNEXt was **Intensity Non-Overlap**, which prioritises RoIs which have little area overlapping with previously fragmented RoIs or which have significant intensity increases over those they do overlap with. Intensity Non-Overlap combined with SmartRoI was shown, compared to TopN Exclusion, to increase both fragmentation coverage and “intensity coverage”, the average precursor intensity each peak was targeted at. A fragmentation spectrum from a higher intensity precursor typically has more parent ions from which we can produce higher quality, more interpretable spectra.

An alternative approach to improving comprehensiveness is to discard the real-time feedback loop occurring scan-by-scan. In these “pre-scheduled” methods, a schedule is computed on some representative data, e.g. an all-MS1 “fullscan”. Because this is not done in real-time like DDA, more involved calculations can be performed, and the fragmentation strategy can employ reasoning based on events that have yet to occur in the current MS run but which have been observed in the representative data.

However, in practice, run-to-run variability may compromise these methods — they are not as robust as DDA or DIA. **DsDA** (Dataset-Dependent Acquisition) [3] processes all previous runs in an experiment to plan for the next run — in order to start this sequence it first uses a TopN run. XCMS [22] is used on previous runs to identify chromatographic peaks, and each is assigned a prioritisation score. A high count of appearances and a high maximum observed MS1 intensity increase the score, and a high precursor intensity at any previous MS2 decreases the score. Compared to TopN, DsDA was shown to increase sample coverage and distribute excess MS2 scans to peaks with lower precursor intensity, for which additional fragmentation spectra may aid annotation most.

In this work we present a new approach to pre-scheduled acquisition using a workflow built on maximum bipartite matching algorithms. The maximum bipartite matching problem is a well-studied problem in computing science, and mapping a real-world problem like LC-MS/MS data acquisition to it allows leveraging this existing field of study. The advantage of this approach in practical terms is that a standard algorithm can be used to efficiently compute an optimal schedule for an input set of peaks (a “target list”) allowing limited resources to be allocated to allow more comprehensive acquisitions. This approach was previously used to compute a theoretical upper bound on the number of unique peaks a DDA or pre-scheduled strategy could target in a single LC-MS/MS run [4]. We have taken this benchmarking tool and developed a practical workflow which can be used to schedule acquisitions. This workflow can be used flexibly to generate a completely pre-scheduled acquisition, or can generate inclusion windows to augment a DDA method.

Additionally, the original matching method had some notable limitations. We therefore have made three main algorithmic improvements to it. First we allow it to compute a global schedule across multiple LC-MS/MS runs (and multiple samples) allowing comprehensive targeting. Secondly, we use a maximum weighted bipartite matching algorithm to optimise the precursor intensity of each target, where the previous technique did not consider the quality of individual assignments. Thirdly, we have extended it so that leftover scans can be redundantly assigned to improve the robustness of the technique. Through computational experiments we will show that the pre-scheduled version of this workflow in theory offers highly comprehensive acquisitions with a minimal number of LC-MS/MS runs. We will also investigate the trade-off between pre-scheduled and DDA methods in robustness, and show that the pre-scheduled method nonetheless has state-of-the-art performance even when exposed to inter-sample variation. In future, this offers a potential roadmap to methods which are highly comprehensive, interpretable and robust.

## 2 Algorithms

To optimally assign MS2 scans to targets by bipartite matching, we model scans and target peaks as a bipartite graph. A bipartite graph is an abstract formalism allowing us to capture relationships between two disjoint sets of objects. Objects are represented as “vertices”, and relationships as an “edge” joining two vertices. A maximum matching is the largest set of pairs of those objects where none of those objects appear in more than one pair. In our scan-peak graph, the presence of an edge represents the ability of an MS2 scan to target a chromatographic peak. Thus solving for the maximum bipartite matching on this graph effectively gives a list of MS2 targets to use for expected maximum performance. The mapping of this problem to a well-understood graph theory problem [8, 17] allows us to solve it efficiently with a standard algorithm (in this case the Hopcroft-Karp algorithm [11]).

In this scan-peak scheme, a bipartite graph can be created from a scan schedule, a target list and a list of representative fullscan .mzMLs, one per LC-MS/MS run to plan for. The scan schedule is a list of RTs and MS-levels for scans to be run, and the target list specifies acceptable (*rt, m/z*) bounds for each target you would like to acquire. The MS1 values in the representative data are used to populate the expected MS1 intensities of each target in the new scan schedule — this is necessary to populate the edges of our scan-peak graph. The maximum matching computed on this graph can then be used directly to create a pre-scheduled strategy, or converted to inclusion windows (by taking a small window around each planned MS2) and given to a compatible DDA strategy. An overview can be seen in Figure 1.

**Figure 1:**
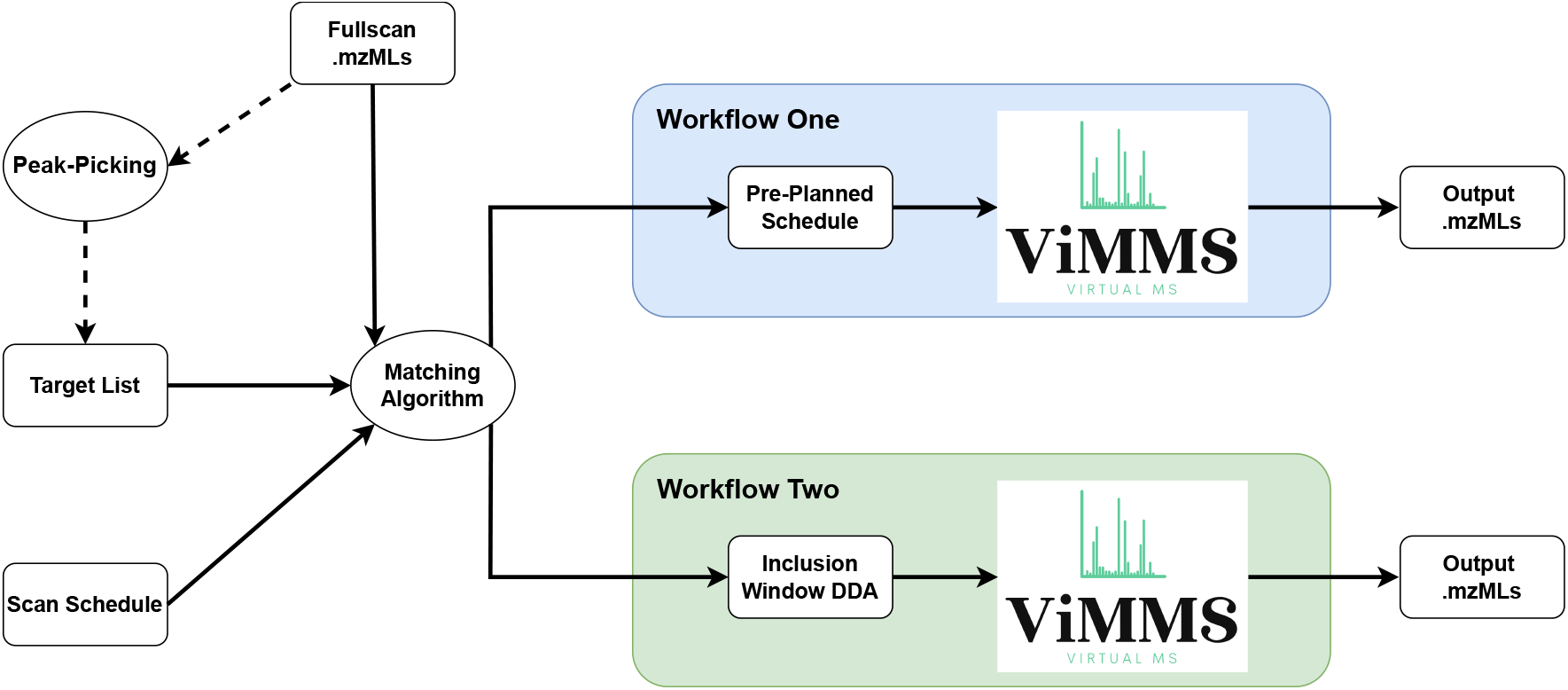
Diagram showing the two possible workflows and the flow of data. In either case, representative (fullscan) data, a target list and a scan-schedule are used to create a maximum matching. This maximum matching can then be converted into either a completely pre-planned schedule, or inclusion windows to inform a DDA method. These can then be run through the Virtual Metabolomics Mass Spectrometer (ViMMS) to produce output LC-MS/MS data. The target list can be produced by any means desired — to produce them in this work we use chromatographic peak-picking software on the representative data. The optional nature of this procedure is represented with dashed arrows.

In Section 2.1 we describe the background of the scan-peak bipartite matching technique as it was previously used [4]. We contribute the ability to actually execute this schedule in an MS run and several improvements described in the following sections. Section 2.2 describes how a scan-peak graph can be constructed for an experiment of multiple MS runs using either a single sample or multiple samples. Section 2.3 describes how we assign MS2 scans to the highest expected precursor intensity available per peak and therefore optimise acquisition quality. Section 2.4 describes how we assign leftover MS2 scans into a full assignment for redundancy and therefore fault-tolerance. Finally, Section 2.5 describes how inclusion windows can be integrated into the TopNEXt framework, and therefore how maximum matching can be used to improve its DDA controllers.

### 2.1 Background

Formally, a graph *G* has vertices *V* and edges *E*. An (undirected) edge is of the form {*u, v*} where *u, v* ∈ *V*. Each vertex represents an object of interest, and an (undirected) edge represents a symmetric relationship between them. In our case we want to capture two disjoint sets of objects, MS2 scans and chromatographic peaks, and model the relationship of whether a given scan is able to target a given peak.

As before, an MS2 scan is represented by a retention time at which it occurs, and a chromatographic peak has a bounding interval in (*rt, m/z*) space i.e. a rectangle. MS2 scans follow an arbitrary schedule given as input, and chromatographic peaks may be provided by XCMS [22], MZMine [18] or an arbitrary method of the experimenter’s choice. We therefore have two disjoint sets of vertices that are subsets of *V* i.e. *S* ⊂ *V, P* ⊂ *V* and *S* ∩ *P* = ∅ where *S* is the set of MS2 scans to be assigned and *P* is the set of target peaks. An edge is created between a given *s* ∈ *S* and *p* ∈ *P* only if the RT of *s* intersects the peak interval and there exists a valid precursor (represented as a single point on the precursor MS1 scan) above a minimum intensity threshold inside the peak interval.

Because we have two disjoint sets of vertices *S* and *P*, and each edge connects a vertex *s* ∈ *S* to a vertex *p* ∈ *P, G* is a bipartite graph. We now want to compute an assignment of peaks to scans such that each scan is assigned to no more than one peak (so that each ion species is isolated individually) and so that scans are assigned to as many unique peaks as possible. A matching on a bipartite graph is a subset of edges such that no two edges in the matching share a vertex i.e. *e*_1_ = {*s*_1_, *p*_1_} ∈ *M, e*_2_ = {*s*_2_, *p*_2_} ∈ *M* ⇒ *s*_1_ ≠ *s*_2_, *p*_1_ ≠ *p*_2_ for a matching *M* ⊂ *E*. A graph’s maximum matching is a matching on that graph such that no larger matching exists on it. The Hopcroft-Karp algorithm [11] is able to solve the maximum bipartite matching problem in 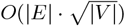 time — we use an implementation provided by NetworkX [10]. Figure 2 illustrates a toy example of the maximum bipartite matching on a graph constructed this way.

**Figure 2:**
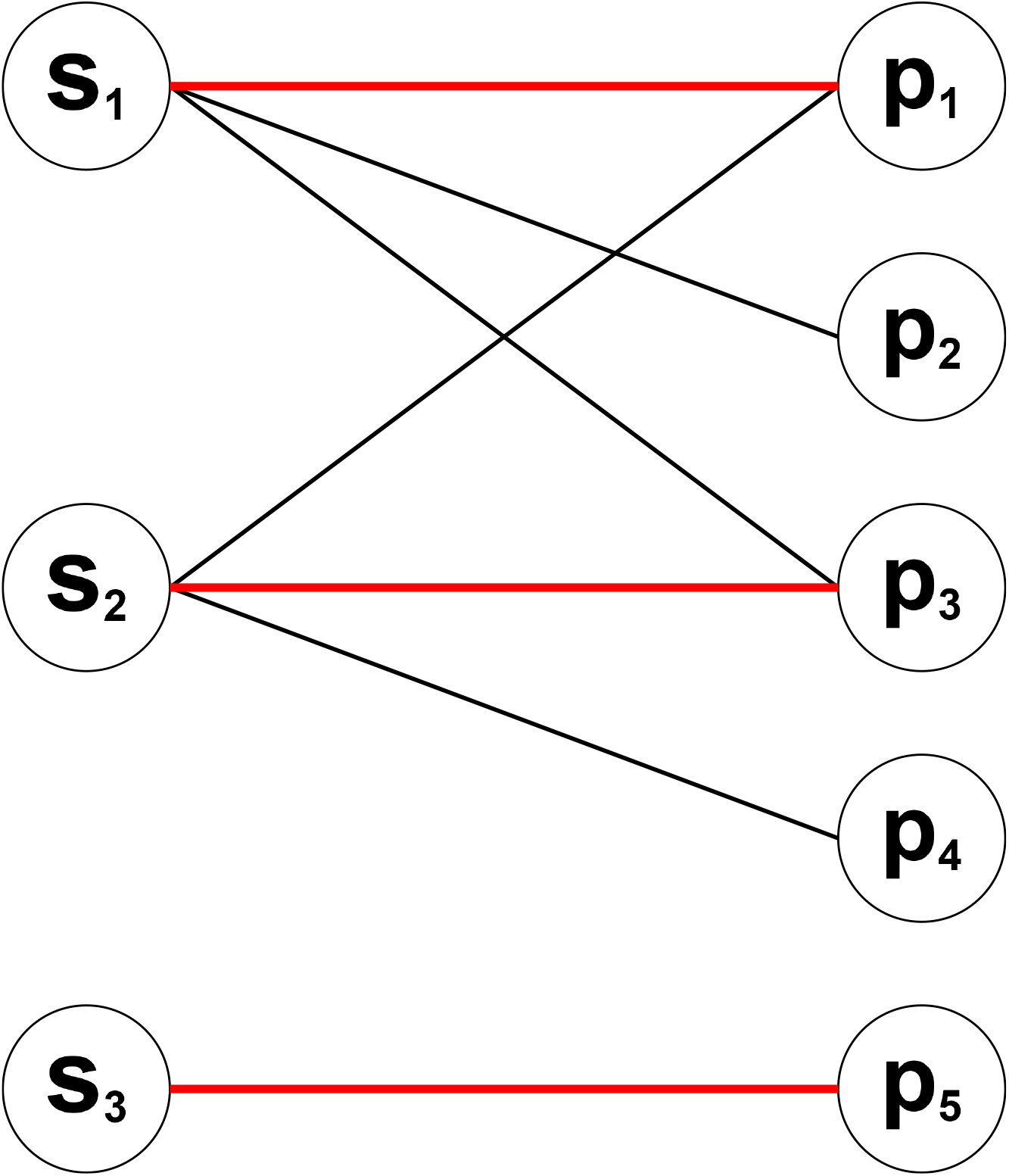
A toy example of a maximum bipartite matching between scans and peaks, where an edge indicates that a peak can be targeted by its connected scan. Edges included in the matching are marked in red and are slightly thicker. All the vertices in the left-hand (scan) side of the matching are assigned — it is “one-sided perfect” or “full”. An obvious consequence of this is that it must also be a maximum matching.

### 2.2 Multi-Sample Matching

In Figure 2 we illustrated planning for a single LC-MS/MS run, but often we may be interested in planning for a series of multiple aligned LC-MS/MS runs (potentially of multiple sample types) to maximise coverage for them as a set. Supposing we create a scan/peak graph for each run, these graphs would likely not be entirely independent. While the scans in each graph would be disjoint from any other graph (they represent different scan events happening across different LC-MS/MS runs) the aligned peaks would not be (if an analyte produces the same peak across two runs we may not want to acquire it twice).

A straightforward, greedy approach is to run the matching algorithm, delete any peaks you successfully targeted, and then run a new matching on the remainder. But this technique is not globally optimal. Consider *p*_3_ in Figure 3A. Should *s*_2_ be naively targeted at *p*_1_ or *p*_4_ instead, then *p*_3_ cannot be acquired as it disappears in the second run in Figure 3B. A greedy method has no foreknowledge that *p*_3_ is not going to be present in run two and thus may miss it. Assuming this disappearance will also occur in the fragmentation runs, then this should be avoidable. An example of a real-world scenario where an event like this might occur could be that the two graphs were generated from a case-control setup where *p*_3_ represents a meaningful biomarker only present in one sample type.

**Figure 3:**
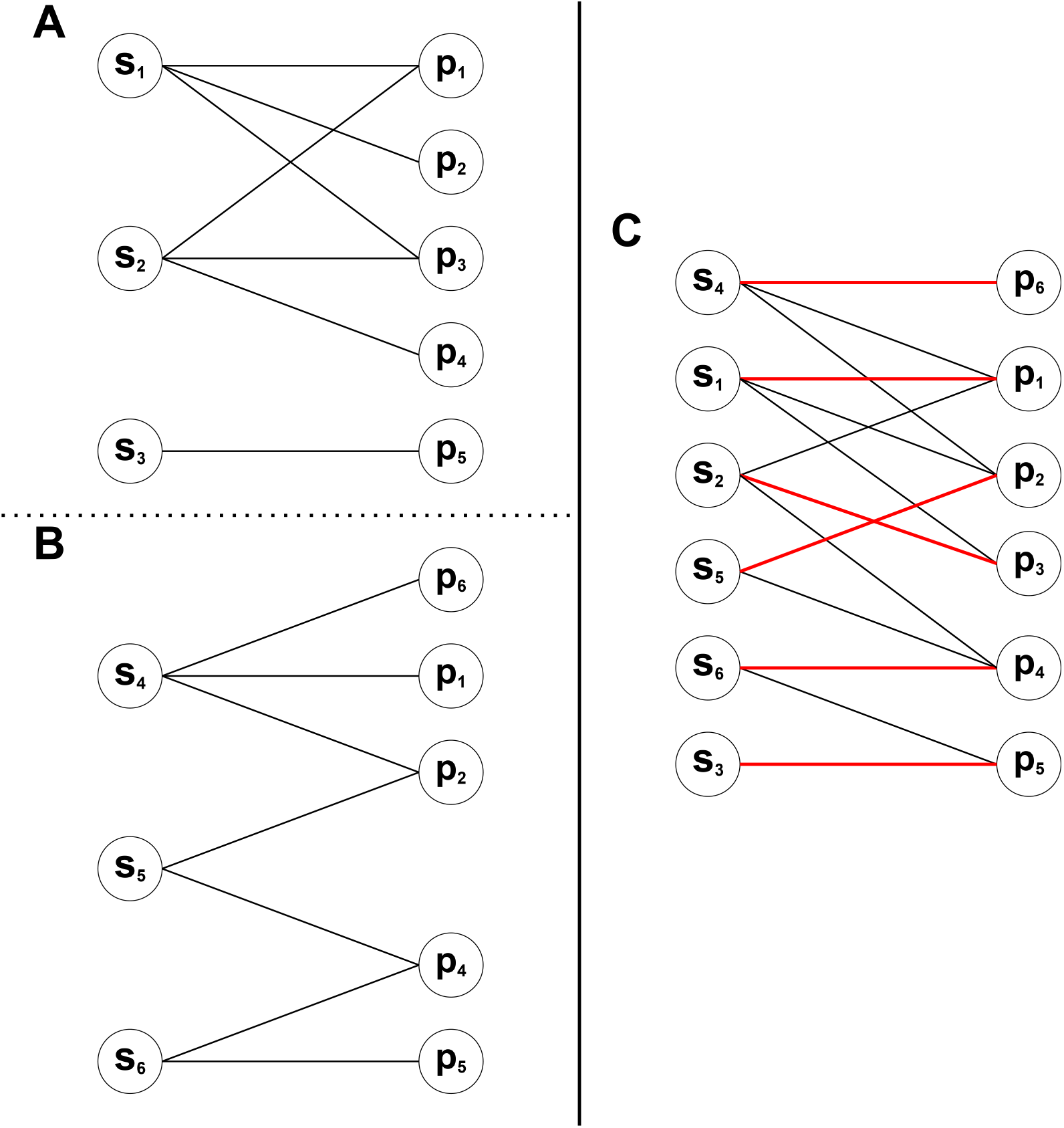
**A** and **B**: Individual graphs for a toy example of two runs of a mass spectrometer. **A** is from Figure 2. **B** is mostly similar, but *p*_6_ appears in *s*_4_, *p*_2_ appears in *s*_5_ instead of *p*_1_, *p*_3_ does not appear at all and *p*_4_ appears in *s*_6_. **C:** The combined version of the two previous graphs for a multi-sample matching. In order to combine the graphs, the scans are “stacked” and the peaks are “merged”. All 3 + 3 = 6 scans appear in the final graph, but each unique peak appears only once. All scans and peaks have been assigned, so this matching is “perfect”. Note that in the combined graph scans have been reordered to reduce visual clutter only.

We therefore instead combine the graphs into a single graph: all scans and edges are preserved as-is, and all *unique* peaks (as decided by peak alignment in e.g. XCMS) are also included. The combined graph in Figure 3C assigns all six peaks because (having observed the representative data) it has foreknowledge that *p*_3_ must be acquired on the first LC-MS/MS run. Conversely, a greedy approach does not look for cases like this and thus will only obtain the optimal solution by luck some of the time. There are caveats to solving a matching on a combined graph, however. The number of runs must be specified in advance, and there is no priority given to acquiring a peak sooner rather than later.

Consider *p*_5_ in Figure 3. If *s*_6_ was not able to target *p*_4_ then the only target available to either *s*_3_ and *s*_6_ would have been *p*_5_, and the matching solver would be indifferent to the resultant choice between *s*_3_ and *s*_6_. The solver might choose to target *p*_5_ with *s*_6_, but suppose that in the fragmentation runs the edge {*s*_6_, *p*_5_} disappeared while {*s*_3_, *p*_5_} remained. *s*_3_ would have already occurred by the time *s*_6_ was observed, so despite *s*_3_ being unassigned there would be no opportunity to target *p*_5_. A greedy approach might try *p*_5_ on the first run, then the second, and so on until it succeeds. If a peak is deferred until a later scan one acquisition opportunity is lost, which may be problematic due to noise.

To address these issues, firstly, the size of the matching can be seen prior to running it, allowing the number of LC-MS/MS runs needed to be estimated in advance. Secondly, it is possible to combine the greedy and combined graph approaches by creating a combined graph for batches of runs — we leave the details of this implementation and selection of chunk size to future work. Thirdly, creating a full assignment as in Section 2.4 allocates scans redundantly to help avoid this issue.

### 2.3 Intensity Matching

As we have mentioned, it is generally preferable to target peaks at their apex precursor intensity to obtain the highest-quality fragmentation spectrum. However, a regular maximum bipartite matching will ensure as many peaks as possible have an MS2 scan assigned, but the algorithm is completely indifferent as to which scan is used provided the choice does not block other targets. Thus, to obtain higher-quality fragmentation spectra, we annotate each edge with the precursor intensity and solve a maximum *weighted* bipartite matching.

However, it is common for algorithms and their implementations to assume that a “perfect” or “onesided perfect” solution is available [19]. The algorithm implemented by NetworkX [13] requires the solution to be “full” (i.e. one-sided perfect). In practice this means if there is no solution where either all scans or all peaks can be included, the algorithm is not valid and cannot be used (the implementation will terminate without finding a solution if it finds this condition is not met). NetworkX does contain an implementation of a standard algorithm for a maximum weighted matching on general (i.e. not necessarily bipartite) graphs [6, 8] but we found that this algorithm was computationally infeasible for our scan-peak graphs.

Therefore to have a working algorithm with an acceptable runtime, we use a “two-step matching” approach. We firstly find a maximum matching, creating an auxiliary graph by deleting any peaks not included in the matching and any edges attached to them. We then secondly solve a one-sided perfect matching on the auxiliary graph using the NetworkX implementation of the Jonker–Volgenant algorithm [12]. Figure 4 shows how the targets might be “swapped” from the initial assignment in Figure 2. This has the side-effect that even if it is otherwise possible to increase the total sum of acquisition intensities by reducing the number of unique peaks targeted, any matching we create will first prioritise the number of unique peaks targeted.

**Figure 4:**
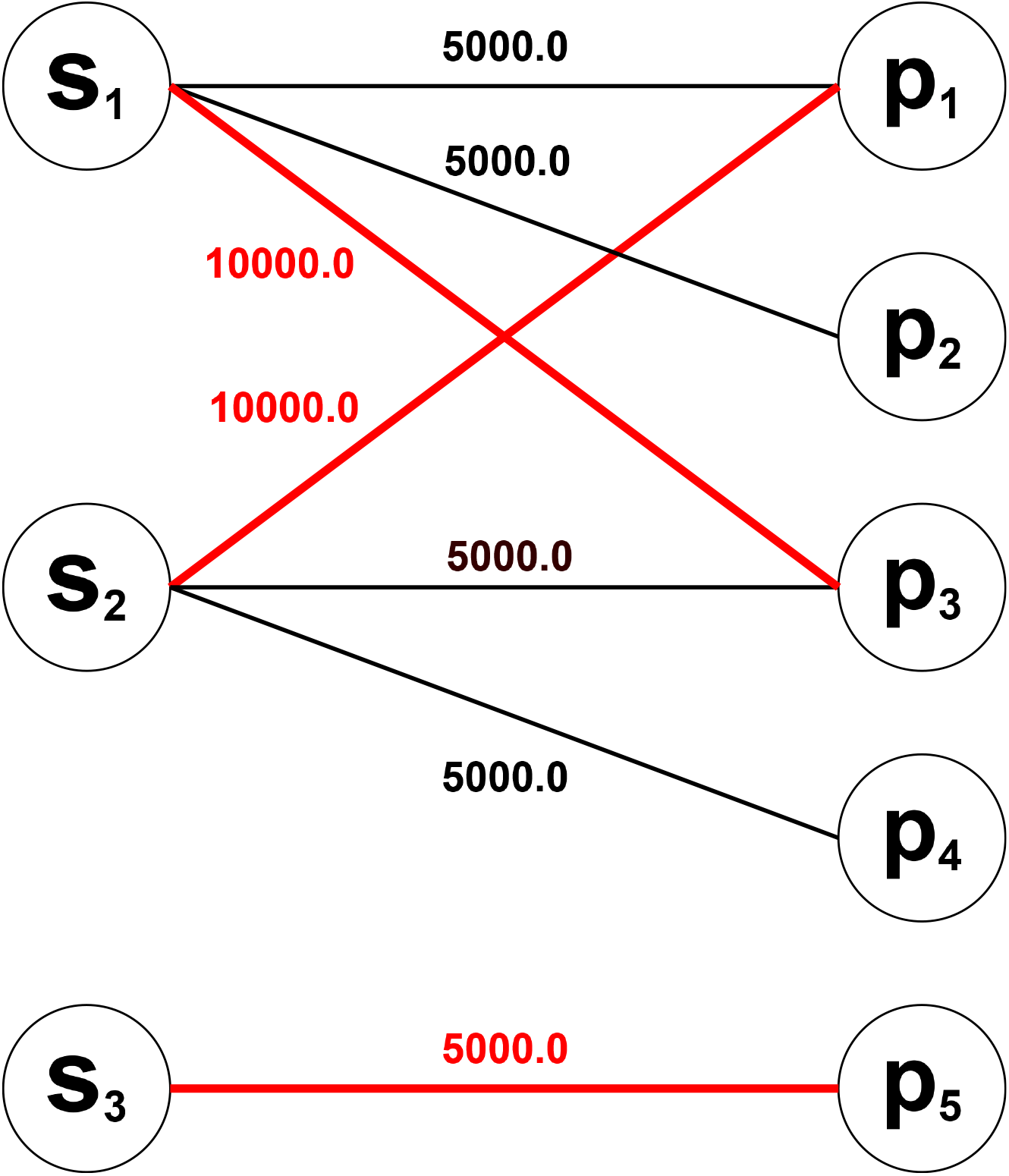
A maximum intensity matching on the toy graph from Figure 2 after precursor intensities have been annotated on the edges. In the two-step matching, we first perform the maximum coverage matching. Assuming we get the same matching as in Figure 2, *p*_2_ and *p*_4_ will not be included in it and thus will be removed to create the auxiliary graph. However, because there is a higher intensity edge from *s*_1_ to *p*_3_ and *s*_2_ to *p*_1_ these will be reassigned and the final matching will have a different assignment of edges to the matching in Figure 2. Any edge between a scan-peak pair is maintained.

### 2.4 Full Assignment of MS2 Scans

As we use more LC-MS/MS runs (for example, to profile crowded regions) we will inevitably end up in a situation where not every MS2 scan can be given a unique target. For example, Figure 5 shows a simple case where there are more scans than peaks. By the definition of a bipartite matching, not all of these scans can be included, but they should still ideally be given a useful target. For example, they may redundantly target peaks in the matching to add robustness or aid identification with extra spectra.

**Figure 5:**
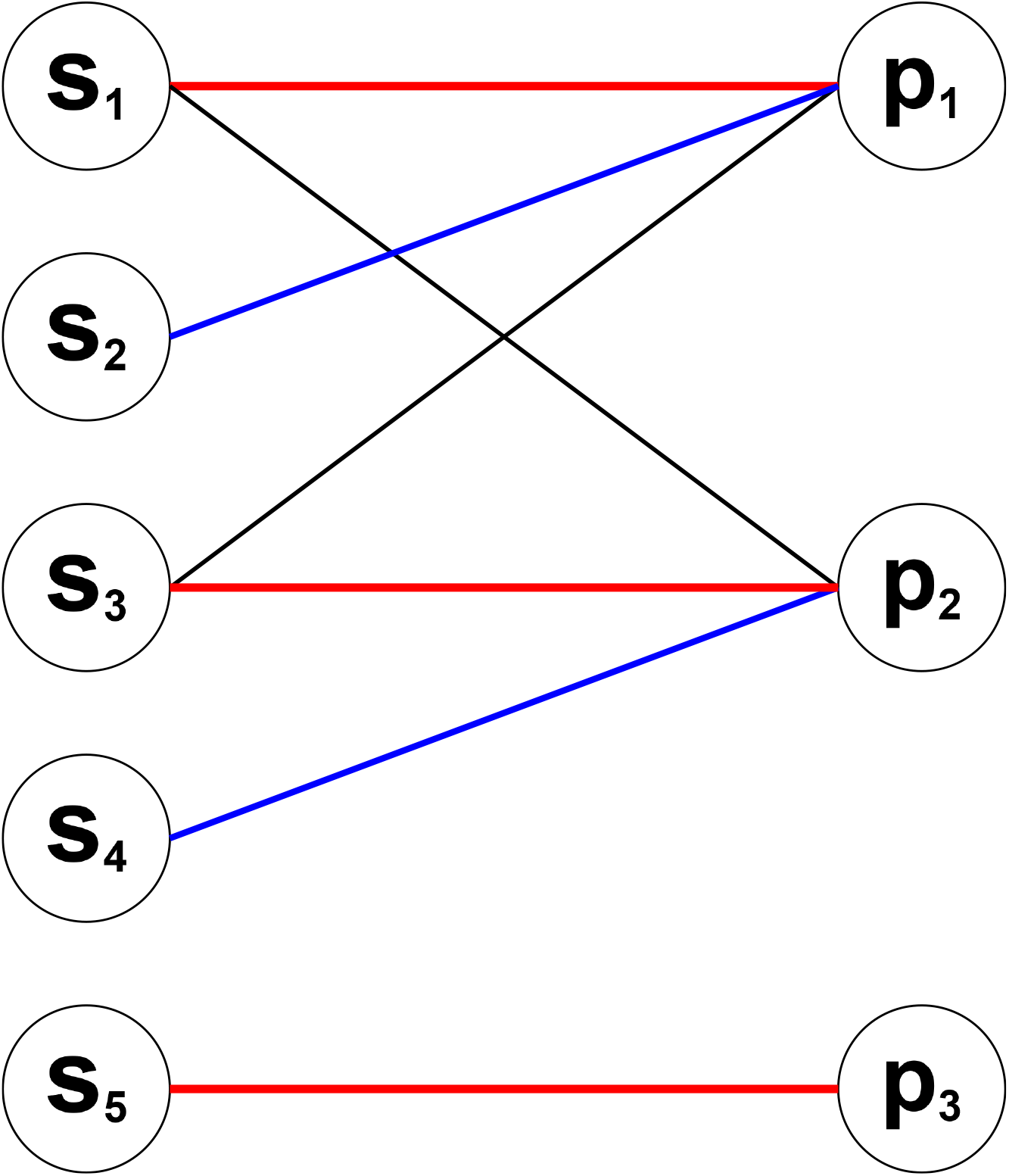
A toy example of a bipartite matching being turned into a full assignment. The graph is the same example given in Figure 2 but flipped so there are more scans than peaks. The matching, marked in red, is also the same but now not all scans are assigned to it. Therefore, marked in blue with lines of less thickness (but more than the unassigned, black lines) is a possible assignment created by, for example, running a second iteration of the matching with *s*_1_, *s*_3_, *s*_5_ removed.

We provide two simple heuristic rules to create a full assignment of scans (though the approach could be easily adapted to use another). The first rule, *nearest*, simply sets any unallocated MS2 scan to have the same target as the nearest (by scan index) allocated MS2 scan (even if the newly allocated MS2 scan did not have an edge to that peak in the matching and thus would not be considered targetable at that point). The second rule, *recursive*, computes a matching, creates an auxiliary graph by removing all peaks not in the matching and all scans in the matching, and then repeats solving on auxiliary graphs and removing scans until no scans remain in the auxiliary graph.

*nearest* is simpler to compute and attempts to redundantly target peaks in a local time frame. *recursive* is much more expensive to compute, taking up to 30 minutes to compute for the experiments in Section 4 (see Supplementary Section 5 for timings). On the other hand, *nearest* has a sub-second timing. In exchange for its more intensive processing, *recursive* has a much better spread on its targets — it attempts to target each reachable peak once for each sub-iteration. This should make it more robust especially against forms of heavy noise such as peaks entirely dropping out (and we will show evidence for this in our experiments). While the computational cost had a reasonable upper bound for our data there may be larger datasets for which a less intensive scheme like *nearest* is required.

### 2.5 TopNEXt Inclusion Windows

Inclusion windows can be straightforwardly generated from a completed matching by creating a window of the desired size around each planned target. These can then be given to a standard inclusion workflow if desired. So that this kind of workflow can be used with advanced DDA methods, we have extended our existing TopNEXt framework [16]. As a reminder, in TopNEXt, a DDA method is expressed as a scoring function of the form in Equation 1.

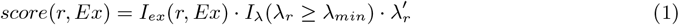

When deciding fragmentation targets, a TopNEXt method assigns each active Region of Interest (RoI), *r*, an initial score 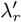 usually based on current intensity. This score is then subject to multiple exclusion criteria. *I*_*λ*_ is an indicator function which ensures the current intensity of the RoI, *λ*_*r*_, is above a user-defined minimum threshold, *λ*_*min*_. *I*_*ex*_ is an indicator function which ensures the RoI is not currently excluded by DEWs or static exclusion windows, which are held in *Ex*. If either of these conditions are not met, the score, given by *score*(*r, Ex*), becomes zero. Individual methods are implemented by defining 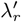. For example, TopN and TopN Exclusion can be implemented by setting 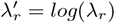, shown in Equation 2.

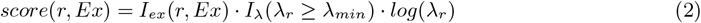

The nominal score for these functions is simply the logarithm of the precursor intensity and the difference between TopN and TopN Exclusion is handled by the exclusion windows stored in *Ex*.

To add inclusion windows to the TopNEXt framework, we included an additional term to the scoring function, seen in Equation 3.

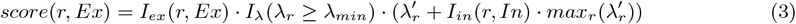

Similarly to *I*_*ex*_, *I*_*in*_ tells us whether or not the RoI falls within any of the inclusion windows in *In*.

If it does then the value of the largest score, *max*_*r*_(*λ*^*′*^_*r*_), is added to the score. This ensures that any valid RoI with the latest precursor falling within an inclusion window will override any RoIs where this condition is not met. However, prioritisation is retained between those RoIs which either both trigger an inclusion window or both do not. This will also not override exclusion windows and the minimum intensity threshold.

## 3 Methods

Our experiments are conducted using the Virtual Metabolomics Mass Spectrometer (ViMMS) [29]. ViMMS contains a digital twin of a mass spectrometer, which can perform realistic experiments on either user-defined “synthetic” chemical samples [28, 30] or a high-fidelity re-simulation of a previous LC-MS/MS experiment [4, 30, 16]. Additionally, the same Python-based code can control a real instrument given it has a compatible API and appropriate bridging code exists between ViMMS and the API [4, 30, 16]. Previous work has shown re-simulations in ViMMS to be highly similar to lab-generated data in a variety of contexts.

For our experiments we use ViMMS’ re-simulation functionality. Depending on what the user wants to achieve, this simulation can be seeded with various kinds of data. In our case, we are going to “replay” an existing lab experiment as accurately as possible under a new scan schedule with accurate simulation of MS1 values. Thus to seed the simulation, we give it one fullscan .mzML per run we wish to simulate (repeats of the same fullscan for multiple runs are allowed). To produce our series of fragmentation .mzMLs under the new scan schedule, ViMMS will interpolate MS1 values between scans (which is why it is important to use a fullscan, for a high MS1 scan rate and thus more accurate re-simulation).

The experiments presented in Section 4 will broadly mimic the experiments we previously performed with TopNEXt, using the same set of samples and similar structure. However, because this time we will be asking how *robust* each fragmentation strategy is, we will be using more fullscans for re-simulation. This necessitated an extra step of data collection, which we will explain in Section 3.1. Additionally, we will be using a similar evaluation procedure to TopNEXt, which we will describe in Section 3.2. However, because this time we are comparing to DsDA, which uses XCMS internally, we will consistently use XCMS for peak-picking to ensure a fair evaluation. Experiments using MZMine are included in Supplementary Section 4. Finally we describe the configuration of fragmentation strategy parameters and XCMS parameters in Section 3.3.

### 3.1 Seed Data Collection

As we have just mentioned, our experiments will differ from those used for TopNEXt [16] by how they use fullscans for re-simulation. Previously, for each type of beer included, we peak-picked one fullscan to produce a target list to evaluate against. If a sample type appeared multiple times in the run sequence, then its fullscan was re-used. Because this time we are interested in investigating the robustness of pre-scheduled methods, we will also include more realistic re-simulation procedures. Because identified peaks can fluctuate quite heavily per-run, we will use a different fullscan (each containing different peaks and fluctuations) for each simulated run. We will also have a separate set of fullscans to generate a target list for pre-scheduled matching methods to plan on, and one set of fullscans for the re-simulation and generating a target list for the evaluation. In total this means for each experiment type (single beer or repeated different beer) we collected 2*n* fullscans, where *n* was the number of re-simulated runs.

When collecting the fullscans to seed the re-simulations, the instrument used for the previous TopNEXt experiments was not available. Seed data was instead collected using a Thermo QExactive Orbitrap mass spectrometer. Negative ionisation mode was used due to positive mode being unavailable — note that this unfortunately lowers the number of analytes successfully ionised and therefore lowers the number of peaks. Fullscan spectra were acquired in negative mode with a resolution of 70,000 and a mass range of 70-1050 *m/z*, and the AGC was set at 1,000,000 for MS1 scans.

The samples used (listed in Supplementary Table 1) were a subset of those used for the TopNEXt experiments, so were retrieved from the freezer. Identically to the data collection procedure for TopNEXt, chromatographic separation with HILIC was performed on all samples by injecting 10 *µ*L beer extract with a Thermo Scientific UltiMate 3000 RSLC liquid chromatography system and a SeQuant ZIC-pHILIC column. A gradient elution was carried out with 20 mM ammonium carbonate (A) and acetonitrile (B), starting at 80% (B) and ending at 20% (B) over a 15 min period, followed by a 2 min wash at 5% (B) and a 9 min re-equilibration at 80% (B). The flow rate was 300 *µ*L/min and the column oven temperature was 40°C. Each beer extract was injected a maximum of 6 times from the same vial before moving to a new aliquot of the same beer extract, in order to minimise over-sampling of the same vial.

### 3.2 Evaluation & Peak-Picking

In this work we evaluate our results in terms of *coverage* and *intensity coverage*. When a strategy successfully targets any MS2 at a unique chromatographic peak above a minimum precursor intensity threshold (5000 in our experiments) said strategy gets one point of coverage. It also receives part of a point of intensity coverage equal to a proportion of the highest precursor intensity the peak was targeted at over the highest MS1 intensity that the peak was observed at. We report these values in proportional form — coverage and intensity coverage are divided by the total number of peaks (counting only those appearing above the same minimum intensity threshold). In this form intensity coverage corresponds to the average percentage of maximum intensity each peak was acquired at. The specifics of how they are calculated can be found in Supplementary Section 3 of the TopNEXt publication [16].

As we have mentioned, running the matching algorithm requires a target list of peaks we would like to target. Similarly, evaluating by coverage and intensity coverage requires a list of desired peaks so we can tell which ones have been targeted successfully. In both cases we use peak-picking on input fullscan (MS1-only) files measuring the same sample as the fragmentation run we want to plan for/evaluate. Specifically, we identify chromatographic peaks on individual fullscans with XCMS’ centwave algorithm [25] and align the peaks with XCMS’ implementation of the MZMine join aligner [18]. Again, the use of fullscans is important as a higher number of MS1 scans in the run will result in higher quality peak-picking.

A peak-picking procedure like XCMS’ typically produces many more chromatographic peaks than there are actual, verifiable chemicals to target (analytes often produce more than one peak, there may be contaminants, etc). But one of our questions of interest is how comprehensive we can make a matching-based method — how many different peaks can it target? Apart from this procedure being automated, the high numbers of targets produced by XCMS are useful in the sense that they make this problem harder. To this end, we give XCMS a particularly liberal set of parameters listed in Supplementary Table 2.

Table 1 illustrates both the high peak numbers produced by XCMS on our data and also how total peak numbers grow as more fullscans are included (hence why we will use a fullscan per run in experiments in Sections 4.2 and 4.3). Note that some peaks do not appear above the minimum intensity threshold within a fragmentation run so will not be counted for evaluation — we report figures in proportional form for this reason. The table also shows peak numbers when using the “Restrictive MZMine” parameter set introduced with TopNEXt. We primarily use XCMS so as not to disadvantage DsDA, but details of the MZMine (Restrictive) parameter set and results using it can be found in Supplementary Section 4. One thing to note is that the total numbers of peaks in Table 1 are lower than an equivalent peak-picking approach run on the datasets generated for TopNEXt — this is most likely a consequence of the instrument being run in negative mode.

**Table 1:** Table showing different output peak numbers for different sets of fullscans being peak-picked and aligned (ranges are inclusive). Fullscans 1-12 are all runs using a single beer sample type. 13-36 use four different beer samples in a repeating pattern. See Supplementary Table 1 for details of the sample types. Note that some of these peaks will be below the required intensity threshold and will not be targeted or counted in the evaluation.

| Fullscan Indices | Num. Peaks<br>(XCMS) | Num. Peaks<br>(MZMine Restrictive) |
| --- | --- | --- |
| 7 | 5784 | 1800 |
| 1-6 | 10385 | 3499 |
| 7-12 | 9671 | 3139 |
| 1-12 | 12626 | 4261 |
| 13-16 | 11804 | 4279 |
| 13-24 | 16916 | 6187 |
| 25-36 | 17741 | 6083 |
| 13-36 | 22293 | 7590 |

### 3.3 Fragmentation Strategy Parameters

Fragmentation strategies often take a number of input parameters, so for our experiments in Section 4 we must select appropriate parameter values. For the strategies we tested previously in our work in TopNEXt (TopN, TopN Exclusion, Intensity Non-Overlap) we will continue using the parameters that were found to be best in that work. These are shown again in Table 2. As we mentioned in Section 3.1, data acquisition was run in negative-only mode due to constraints on instrument availability but the samples themselves have not changed. Although this could alter the optimal choice, the time taken to optimise parameters means it may not be realistic to do it for every experiment. We will also have Intensity Non-Overlap use its SmartRoI variant, because this previously performed best. However, while we use the TopNEXt implementation of TopN Exclusion (“TopNEx”) in order to use matching-generated inclusion windows, we use it with the regular DEW to give an idea of the performance gain when adding it directly to TopN Exclusion.

**Table 2:** Parameters used for DDA fragmentation strategies in our experiments.

| Shared DDA Parameters |  |
| --- | --- |
| Ionisation mode | Negative |
| $N$ | 20 |
| Isolation width | 1 |
| Min MS1 intensity ($\lambda_{min}$) | 5000 |
| $mz\_tol$ | 10 |
| $min\_roi\_length$ | 3 |
| DEW Parameters |  |
| $rt\_tol$ | 60 |
| SmartRoI Parameters |  |
| $rt\_tol$ | 15 |
| $intensity\_increase\_factor$ ($\alpha$) | 3 |
| $drop\_perc$ ($\beta$) | $10^{-3}$ |

Given that it is a state-of-the-art pre-scheduled method, we also include DsDA in our comparisons. Because DsDA uses XCMS internally to iteratively process the previous set of runs in preparation for the next, our main experiments will use XCMS to construct matching and evaluation target lists to ensure a fair comparison with DsDA. DsDA will use the same set of XCMS parameters for its peak-picking as will be used for both target lists (values can be found in Supplementary Table 2). It should be noted that DsDA does not use XCMS’ equivalent of the MZMine join aligner — it groups peaks if their m/z falls within a ppm tolerance and they overlap at all in RT. We have left this as-is in order to minimise the number of changes we made to the method and its implementation.

Other than the XCMS parameters, DsDA has two main parameters to choose values for: *N* and *maxdepth*. So the comparison to the TopNEXt methods would be fair, we chose values by the same optimisation procedure as with TopNEXt i.e. we grid searched values by running re-simulated experiments of the same format later shown in Section 4.1 and choosing the parameter combination that maximised intensity coverage. The *N* parameter controls the number of MS2 scans generated per MS1 scan in the schedule DsDA fills in. A higher value of *N* gives DsDA more opportunities for targeting. But DsDA picks peaks on its previous runs — that means a lower value of *N* means richer information is given to *centwave* and thus DsDA will better be able to pick peaks. A setting of *maxdepth* = *m* causes DsDA to completely exclude any previously-acquired peaks on every *m*th run. This parameter may therefore help if DsDA is prone to missing certain peaks. For *N* we searched the values 1, 5, 10, 20 and for *maxdepth* we searched 1, 2, 3, 4 and NULL (NULL meaning to disable *maxdepth*). For the single-sample experiment we used *N* = 10 and *maxdepth* = 3 and for the round-robin experiment we used *N* = 20 and *maxdepth* = *NULL*.

The matching algorithm used a TopN scan schedule with *N* = 20, the appropriate set of fullscans for each experiment, and the list of peak intervals produced by XCMS when processing and aligning each set of fullscans. Inclusion windows were created with an RT width of 10 seconds (approximately two full duty cycles) and an m/z width of 10 ppm around the target RT and m/z.

## 4 Results

In the previous work on TopNEXt [16], we peak-picked one fullscan per sample type to produce a target list to evaluate against. However, both the underlying data and how a peak-picker will interpret that data can vary significantly per run. To measure the robustness of pre-scheduled methods, we paired each fullscan used for the simulation and evaluation with another “held-out” fullscan that could be used to plan on. Additionally, as Table 1 shows, the total set of peaks increases significantly in number with more runs. To account for this we also use a different fullscan per each run.

We introduce these changes incrementally, so first in Section 4.1 we perform an experiment in the style of TopNEXt where a single fullscan is used for each sample type (so for simulation, evaluation and planning for all runs of that sample type). Unlike the previous experiment, in Section 4.2 we use different fullscans for each run, but we still only use one fullscan to plan, simulate and evaluate each run. Finally in Section 4.3 we collected two equally-sized sets of fullscans, one set to be used for planning, and the other set to be used for simulation and evaluation. For each run we use a different fullscan, but the sequence of sample types is the same. This means that for a given simulated run in this experiment, the planning will use a different fullscan than the simulation and evaluation, but both procedures will share the same underlying sample type. This mimics a scenario where one might collect *n* fullscans to plan on, and then perform an experiment of *n* runs. Of the 2*n* fullscans we collected, the first *n* are used for planning, and the latter *n* for simulation and evaluation. The variability in peaks between runs can be seen in how peak numbers grow as files are aligned in Table 1.

The samples chosen for our experiments are the same as those chosen for TopNEXt, but we run fewer total iterations (because the performance of pre-scheduled methods can be shown in fewer runs). For each choice of how to use fullscans to inform simulation and target list creation, we show both a case of six repeats of a single beer sample, and three repeats of four samples in round-robin order (1-2-3-4-1-2-3-4-1-2-3-4).

Moreover, experiments will consider three classes of method: “baselines” previously investigated by the work on TopNEXt plus DsDA, pure “matching” methods which use the matching workflow to generate a fully pre-scheduled plan and “inclusion window” methods which augment a baseline with the matching workflow by using the matching workflow to generate inclusion windows for the baseline method. Because the inclusion windows are implemented through TopNEXt, the inclusion window version of TopNEXt uses the TopNEXt equivalent of TopN Exclusion but the baseline version does not. Intensity Non-Overlap uses the SmartRoI variant because this performed best in the work on TopNEXt.

Finally, because the matching workflow as we have implemented it knows how many runs it has available in advance and does not prioritise earlier runs, we ran all possible experiments of shorter lengths for both the case of the same beer repeated six times and the different four beers each repeated three times. For the same beer case this meant (1 + 2 + 3 + 4 + 5 + 6 = 21) total runs to produce all experiments of length up to six (and similarly different beers case). We present results for both the run-by-run performance in the experiment of greatest length and the final result for each of these experiments of length up to the maximum. We will be focusing more on the latter case (as it is likely more reflective of how the method would be used in reality) but it is interesting to see how pre-scheduled matching methods allocate scans (particularly when considering the full assignment variants). Furthermore, because there is a possibility of using the inclusion window methods greedily (it is not known prior to running how the DDA component will perform) the run-by-run numbers may be relevant to this use-case.

### 4.1 Re-used Seed Data

We will first consider the case where a single fullscan is used for each sample when generating simulations and target lists. As we mentioned, this experiment is the most similar to the TopNEXt experiments [16], but is not one-to-one comparable due to significantly different numbers of peaks resulting from the seed data being collected in negative ionisation mode and target lists being generated via XCMS rather than MZMine.

Figure 6 shows this re-used fullscan setup for an experiment of the same beer sample being repeated six times. To explain the legend, “INO” is Intensity Non-Overlap (SmartRoI), TopNEX is the TopNEXt implementation of TopN Exclusion (they are two slightly different variations of the same method), TS Matching is the two-step matching with “R” being the variant using *recursive* and “N” being the variant using *nearest*, and a “+” (as in TopNEX + and INO +) denotes the use of matching-generated exclusion windows.

**Figure 6:**
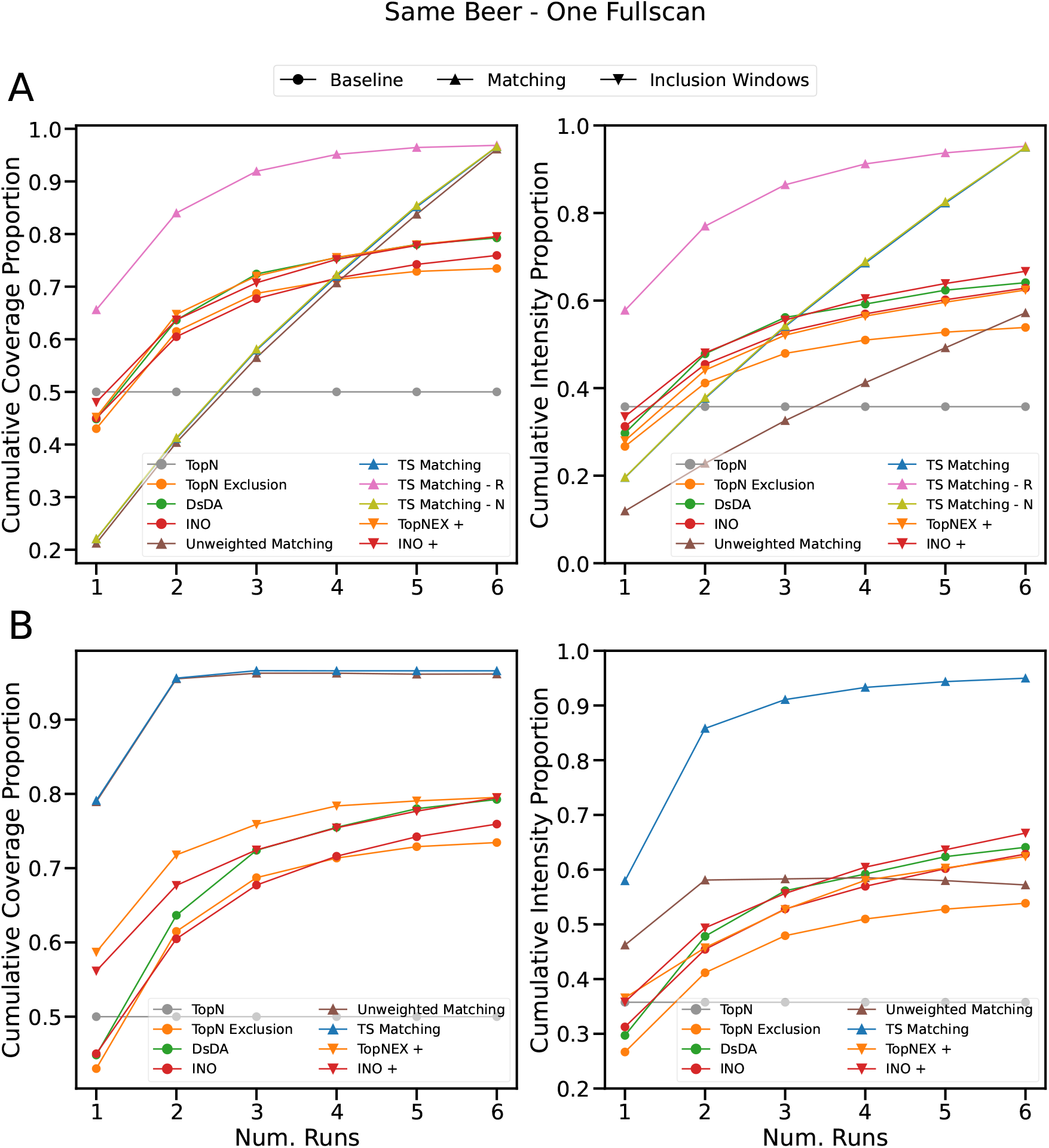
Experiment using the same beer repeated for six samples. A single fullscan was used to generate simulations and target lists, and XCMS was used to generate target lists for both the matching algorithm and the evaluation. **A:** shows performance over different runs in a single experiment. **B:** shows performance over separate experiments of different run numbers.

For the baseline methods studied in our work with TopNEXt the trend is roughly the same. TopN is a flat line well below the performance of those methods which consider information from multiple samples. TopN Exclusion and Intensity Non-Overlap have comparable coverage, and Intensity Non-Overlap has a substantial (approx. 10% of total) advantage in intensity coverage. Coverage caps at around 80% and intensity coverage at around 60% after six runs. This is because of the low level of filtering with our XCMS settings, meaning there are many low-quality targets and making it difficult to obtain total coverage. While we won’t draw close comparisons due to the differing ionisation modes of the two datasets, it should be noted that this is approximately the same final threshold observed using the “permissive” peak-picking parameters in Supplementary Section 5.2 of our work on TopNEXt. The “permissive” MZMine parameters are closer to this XCMS setup in terms of quality threshold than the “restrictive” peak-picking parameters.

Moving on to methods not used in the original TopNEXt study, DsDA is also very competitive, performing best by a small margin in coverage and intensity coverage out of any of the baselines. The pre-scheduled two-step matching method is predictably very strong in this best-case scenario of a noiseless environment where it knows exactly what is coming. The pre-scheduled two-step matching methods outperform all other methods massively, gaining effectively total coverage in an experiment of two runs and over 90% intensity coverage in an experiment of three runs (Figure 6B). Note that the two-step matching is roughly 20% ahead in coverage compared to any other method but is around 30% ahead in intensity coverage, meaning it is even better in terms of intensity coverage. This speaks to the available design space for future methods which attempt to optimise intensity coverage, as improving intensity coverage appears to be relatively unexplored.

We also see the added value of the two-step matching procedure compared to the original matching technique, because while the unweighted matching has close to identical coverage to the two-step matching, it has much lower intensity coverage. It does not outperform the majority of the baselines in intensity coverage despite much higher overall coverage, showing that it acquires spectra at very low average precursor intensity. After the second point on Figure 6B, when it maximises coverage, the intensity coverage approximately flatlines because the unweighted matching is not trying to improve it. There are some minor fluctuations, but this can be attributed to the solver algorithm not making exactly the same choices when presented with a larger graph.

We can also see that the recursive variant of the full assignment distributes its scans much more aggressively over the total number of runs in Figure 6A. Any target arbitrarily chosen for a later run by the matching algorithm may arbitrarily have a duplicate assigned to an earlier run by *recursive*, so the coverage line rises much faster on the run-by-run comparison. Note that because this experiment is noiseless all full assignment variants will have equal performance after the final run, so they have been combined into a single line for Figure 6B.

Finally, the inclusion method windows are a strict improvement on their respective baseline methods. This improvement is especially pronounced at lower levels of runs, where the inclusion does more to help guide the behaviour. With more runs, the iterative exclusion behaviour of the regular baselines allows them to close the gap somewhat. Interestingly, although inclusion windows are still a net benefit to Intensity Non-Overlap, the performance increase is minor as the number of runs increases. Conversely, without inclusion windows TopN Exclusion ends with the weakest coverage of the baselines and a severely lagging intensity coverage. But when inclusion windows are added its coverage is the best of the cluster of non-pre-scheduled methods and its intensity coverage is competitive. The likely implication is that inclusion windows are more of a complement to TopN Exclusion’s inflexible hard exclusion behaviour, whereas while they add information to Intensity Non-Overlap they do not interface as well with its more complex behaviour.

Figure 7 shows the same setup where fullscans are allowed to be reused, but for the case where four beers are repeated three times each i.e. there are four fullscans used three times each by both the simulator and the matching algorithm. Again, similar results to the original tests of TopNEXt can be observed: TopN flatlines once it has seen all the samples, and TopN Exclusion seems to perform comparatively poorly on different-samples cases, especially relative to Intensity Non-Overlap.

**Figure 7:**
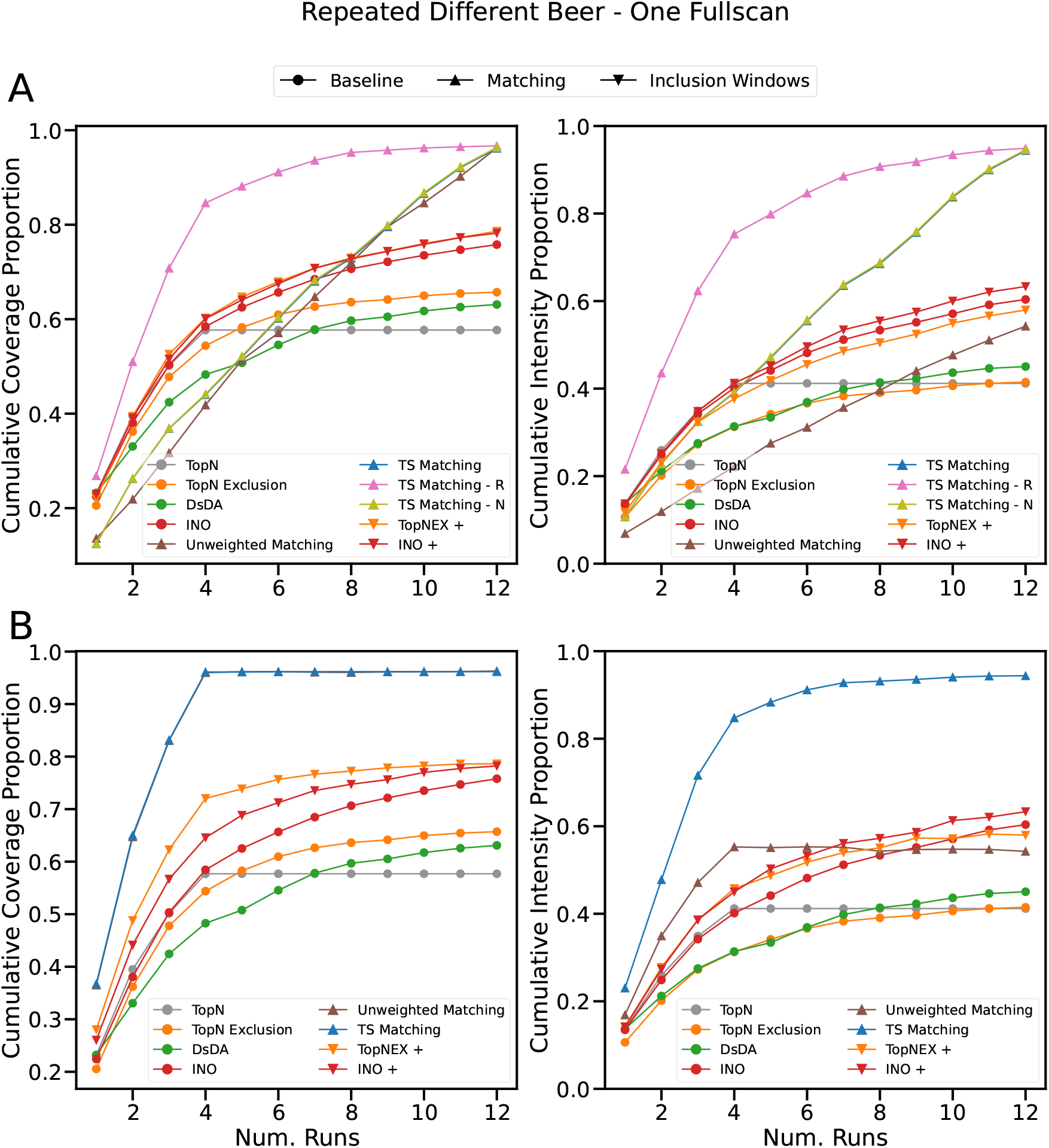
Experiment using four beers repeated three times each in round-robin order (1-2-3-4-1-2…). A single fullscan was used to generate simulations and target lists, and XCMS was used to generate target lists for both the matching algorithm and the evaluation. **A:** shows performance over different runs in a single experiment. **B:** shows performance over separate experiments of different run numbers.

It should be noted that the final thresholds they reach (60 to 80%) are more similar to the experiments with permissive parameters in TopNEXt’s Supplementary Section 5.2, but in that case the relative performance advantage of Intensity Non-Overlap narrowed. The performance advantage for Intensity Non-Overlap was seen in the *main* experiments from the TopNEXt publication. The lack of consistency may be the result of using parameters more similar to the “permissive” parameters, but on a negative ionisation mode dataset with fewer peaks (again suggesting the data is not directly comparable).

We also see that the pre-scheduled matching can effectively obtain total coverage by only seeing each sample once, and close to 85% intensity coverage in the same span. The same observations can be made regarding how aggressively the recursive variant spreads its scans, and also that the unweighted matching has remarkably poor intensity coverage given its comparable coverage.

For the inclusion window methods, we also see again that their use offers Intensity Non-Overlap a minor improvement. And, again, TopN Exclusion is given a substantial improvement by the use of inclusion windows, but this time, the improvement appears to be substantially stronger (around twice as much coverage gained). This gives increased credence to the idea that inclusion windows help compensate for some of the natural weaknesses of TopN Exclusion, as it also struggled with this experiment structure when being tested against TopNEXt.

One thing that has changed from the single sample example, however, is that DsDA is no longer effective, only barely outperforming TopN (and only once it has seen all the samples twice). This makes some sense: DsDA is by design trying to target commonalities in the sample set, and which common peaks are observed will grow as more runs are observed and shrink as the underlying samples get more biologically diverse. DsDA may therefore require many runs (the original work tested it with 20, 40 and 50 [3]) and it may be the case that above a certain level of sample diversity in a multi-sample experiment DsDA will fail to converge to collecting the non-shared parts. If it is the case that DsDA will hit a fundamental limit at a certain level of sample diversity, then in practice it will be necessary to perform data acquisition with DsDA for each sample separately.

### 4.2 Per-run Seed Data

Now we will move onto a different kind of experiment from the ones used for testing TopNEXt. Figure 8 shows an experiment with the same beer repeated, but with a different fullscan being used for each run. Each of these fullscans is used for simulation and evaluation, and planning by the matching algorithm. The first obvious change from the experiments we have seen previously is that TopN is no longer a flat line: the increased number of fullscans being used causes there to be a greater number of peaks only in a certain subset of scans, so there is always new peaks for TopN to acquire. Interestingly, this also makes TopN much more competitive relative to the other methods. TopN Exclusion and DsDA barely outperform TopN, though Intensity Non-Overlap does maintain a significant advantage.

**Figure 8:**
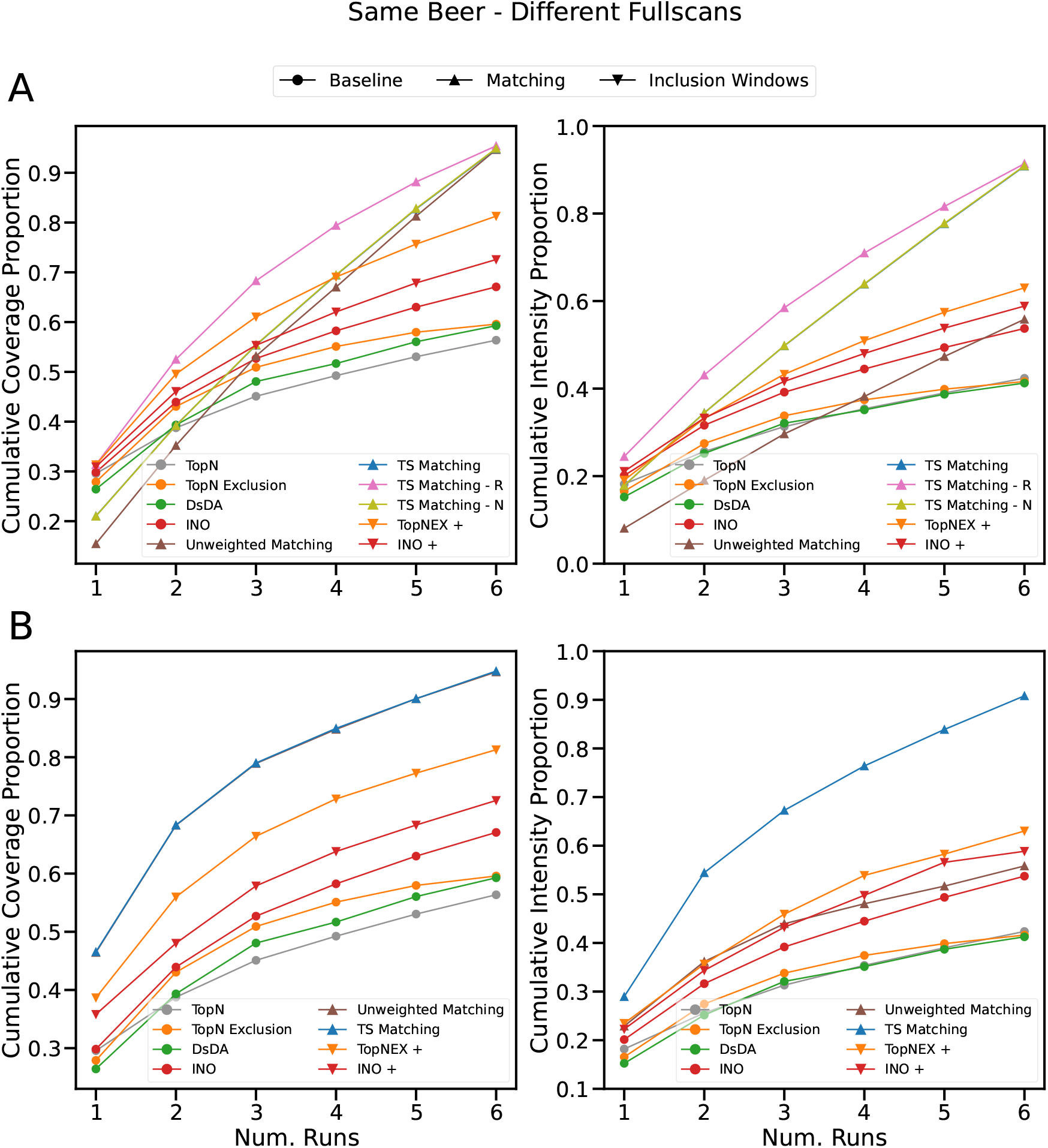
Experiment using the same beer repeated for six samples. A different fullscan for each of the six runs was used to generate simulations and target lists, and XCMS was used to generate target lists for both the matching algorithm and the evaluation. **A:** shows performance over different runs in a single experiment. **B:** shows performance over separate experiments of different run numbers.

The overall high performance of the pre-scheduled two-step matching has not changed, but because more peaks are being introduced per each sample, we do see that it takes longer to reach its apex, and that apex is closer to 90% than before. What is interesting is that the advantage for the baseline methods in having inclusion windows added seems significantly more pronounced (e.g. a 20% coverage increase for TopN Exclusion). It may be the case that having prior information about the distribution of these added peaks provides a significant advantage, but (as we will see in Section 4.3) this advantage cannot be carried forward into reality, given that peaks randomly showing up in one run or another will happen, by definition, in a random order. If, however, it is the case that some of these peaks are below the normal limit of detection given only a single sample, but the alignment of multiple samples allows their detection, then this improvement may partially translate into reality.

Now we will consider the repeated different beers in Figure 9. Surprisingly, TopN actually significantly outperforms both TopN Exclusion and DsDA! In DsDA’s case, this is likely a matter of looking for common peaks to target in an environment where both samples and individual peaks are constantly shifting. For TopN Exclusion, this seems like an exacerbated version of the drop in performance that multiple samples cause. However, it is possible that part of this could be attributed to TopN Exclusion having exclusion windows that are too wide, or to XCMS inappropriately splitting peaks that should be merged together. If this were true it would speak to the well-known difficulties in LC-MS/MS for configuring tools and in comparing results across different configurations. Regardless, Intensity Non-Overlap shows strong performance and is the best-performing baseline.

**Figure 9:**
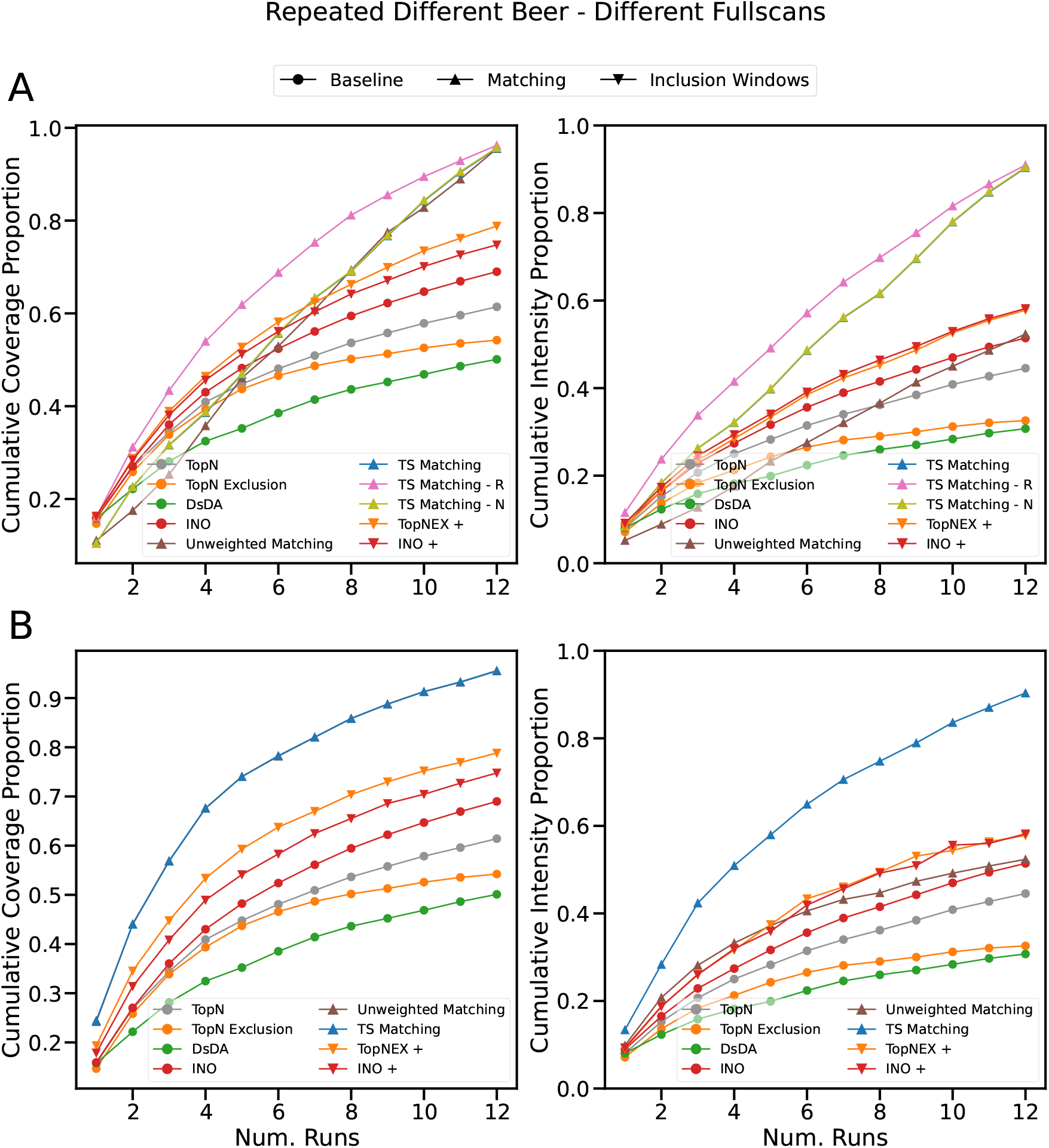
Experiment using four beers repeated three times each in round-robin order (1-2-3-4-1-2…). A different fullscan for each of the twelve runs was used to generate simulations and target lists, and XCMS was used to generate target lists for both the matching algorithm and the evaluation. **A:** shows performance over different runs in a single experiment. **B:** shows performance over separate experiments of different run numbers.

For the pre-scheduled two-step matching, we see the same effect as we did for a single beer type repeated. Performance is very high, but takes a lot longer to accrue and caps at a slightly lower threshold. The trends with the inclusion window methods are also the same as before, other than the fact that the difference between the inclusion window variant of TopN Exclusion and its baseline version has become even larger: again suggesting it has some corrective effect.

### 4.3 Paired Seed Data

So far we have been looking at what is essentially the “best-case scenario” for our matching methods (both in terms of pre-scheduling and inclusion windows) but it is now time to challenge it with a difficult (and more realistic) scenario. In these experiments we will have two sets of fullscans, one to generate a target list for the matching algorithm, and the other to seed the simulator and create a target list for the evaluation. This causes the matching algorithm to see a dataset of the same underlying samples in the same order, but with different peaks, and with peaks showing up in different runs in the run order (as would happen in reality). This is intended to mimic the real-world scenario where one could run an initial experiment, collecting fullscans to plan an acquisition, then run the acquisition afterwards. We have given these two stages the same number of fullscans to account for the fact that peak numbers generally increase as you peak-pick a larger set of fullscans (there is a possibility that this would be too many fullscans to collect in a realistic scenario, but we leave this to the reader’s discretion).

Figure 10 shows the results of this setup on the experiment with the same beer repeated. The baselines were not re-run between this experiment and the previous one, as they do not take a target list as input, so are displayed at the same values as before. Predictably, the performance of the matching-based methods is no longer as good. They are still among the best performing methods, however. Pre-scheduled two-step matching with recursive assignment, and both inclusion window methods have similar performance to Intensity Non-Overlap by the sixth run (with Intensity Non-Overlap plus inclusion windows having the best performance by a slight margin) but accrue performance faster in the earlier runs.

**Figure 10:**
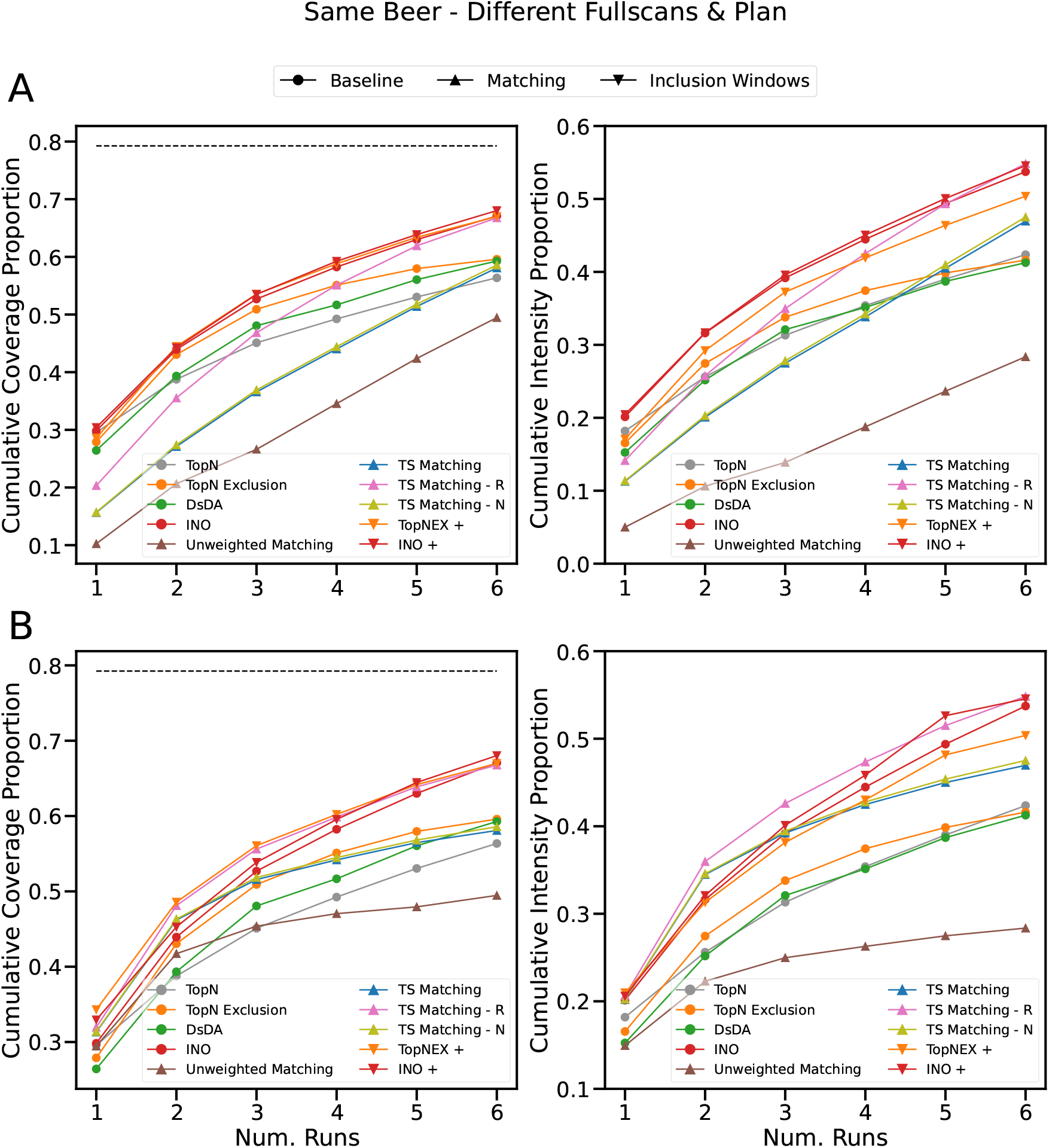
Experiment using the same beer repeated for six samples. Two sets of six fullscans were used. One set was used to generate the target list for the matching algorithm, and the other was used for the simulations and the target list of the evaluation. XCMS was used to generate target lists for both the matching algorithm and the evaluation. The dotted line indicates the level of overlap between the target lists generated from the two sets of fullscans. **A:** shows performance over different runs in a single experiment. **B:** shows performance over separate experiments of different run numbers.

Something to note is that because we have introduced a source of error for the matching workflow, Figure 10B shows all three full assignment variants of the two-step matching. A and B otherwise both show the same content as in previous figures, but now the values between the three variants can differ in B, so they have been plotted separately. Notably, the way the recursive variant spreads targets across scans gives it a significant advantage here — *nearest* barely outperforms the base variant, and they are close to the weaker baselines in coverage after enough runs have passed.

It may be worth observing, however, that while DsDA, TopN and TopN Exclusion catch up to non-recursive variants of the two-step matching in coverage, they still lag behind in intensity coverage (by approx. 5%). Though it is less pronounced, we can also observe in Section 4.2 that these three baselines seem to perform slightly worse in terms of intensity coverage relative to coverage when compared to the other methods. We have seen previously through TopNEXt that this is true for TopN and TopN Exclusion relative to a method like Intensity Non-Overlap, because these baseline methods don’t go back to optimise intensity coverage. Like Intensity Non-Overlap, the matching methods were designed with optimising intensity coverage in mind, so this is evidence that they are effective at this goal.

DsDA, however, was designed with optimising target acquisition in mind — though DsDA’s authors measured this by plotting each time a peak was re-targeted, and found that DsDA “oversampled” low-intensity peaks more than TopN. It seems likely that intensity coverage gives us a slightly different view of this phenomenon by directly showing us the maximum intensity each peak was targeted at. It may be the case that DsDA simply requires more runs to optimise intensity coverage (perhaps with an appropriate setting of the *maxdepth* parameter) or it may be the case that while it may target peaks more often it does not do so at higher intensity or it may be the case that DsDA’s scoring function somehow “saturates” because it is based on the relative rankings of the peaks it sees. These would be interesting questions to answer in future.

Another thing to consider is that the unweighted matching has the worst performance of any method here, being strictly worse than any other method after the third run. One of the most interesting aspects is that while its intensity coverage has always been poor (after all, it pays no attention to targeting intensity) its coverage is also significantly worse than the two-step matching here — this was not true previously. The likely reason is that trying to optimise intensity coverage leads the two-step matching to target the more stable peak apex, where the unweighted matching may arbitrarily select an edge point that will not be in the peak if the peak moves between runs. Together with the observations about the recursive variant, this demonstrates the necessity of our updates to the matching technique for it to be usable as a practical fragmentation strategy.

One final thing to take note of is that the level of agreement between the target lists has been indicated by a dotted line on the plot. In addition to being peak-picked separately, both sets of fullscans were peak-picked together as one set and aligned. The dotted line shows how many of the peaks listed in the matching workflow’s target list were included in the evaluation’s target list. In other words, if a matching method hit every target in its list (and only those targets) it would receive a score of just over 80%. Of course, the actual score for even the best performing pre-scheduled matching is much lower, showing that it is missing a good number of its targets due to those targets, for example, not being in the right run, or right location, or not appearing at all.

Finally we will consider the case where there are two different sets of fullscans for four beers repeated three times each, shown in Figure 11. Surprisingly, under these conditions, TopN is a very effective method, beating the majority of its competitors. This includes the unweighted matching, DsDA, TopN Exclusion, and the regular and *nearest* variants of the two-step matching. It also generally remains the case that inclusion windows are a mild improvement for Intensity Non-Overlap, which finishes as the most effective method, but the pre-scheduled two-step matching using *recursive* gains performance faster and is competitive by the end. TopN Exclusion with inclusion windows is also competitive with both but has generally worse intensity coverage and a slightly lower coverage point by the end. We also see that TopN Exclusion and DsDA do even more poorly in intensity coverage compared to Figure 10 which again suggests they struggle with this multi-sample case. TopN has caught up with the two-step matching methods, but notably the advantage it has (presumably because of the large differences in peaks between runs) is more in coverage than intensity coverage.

**Figure 11:**
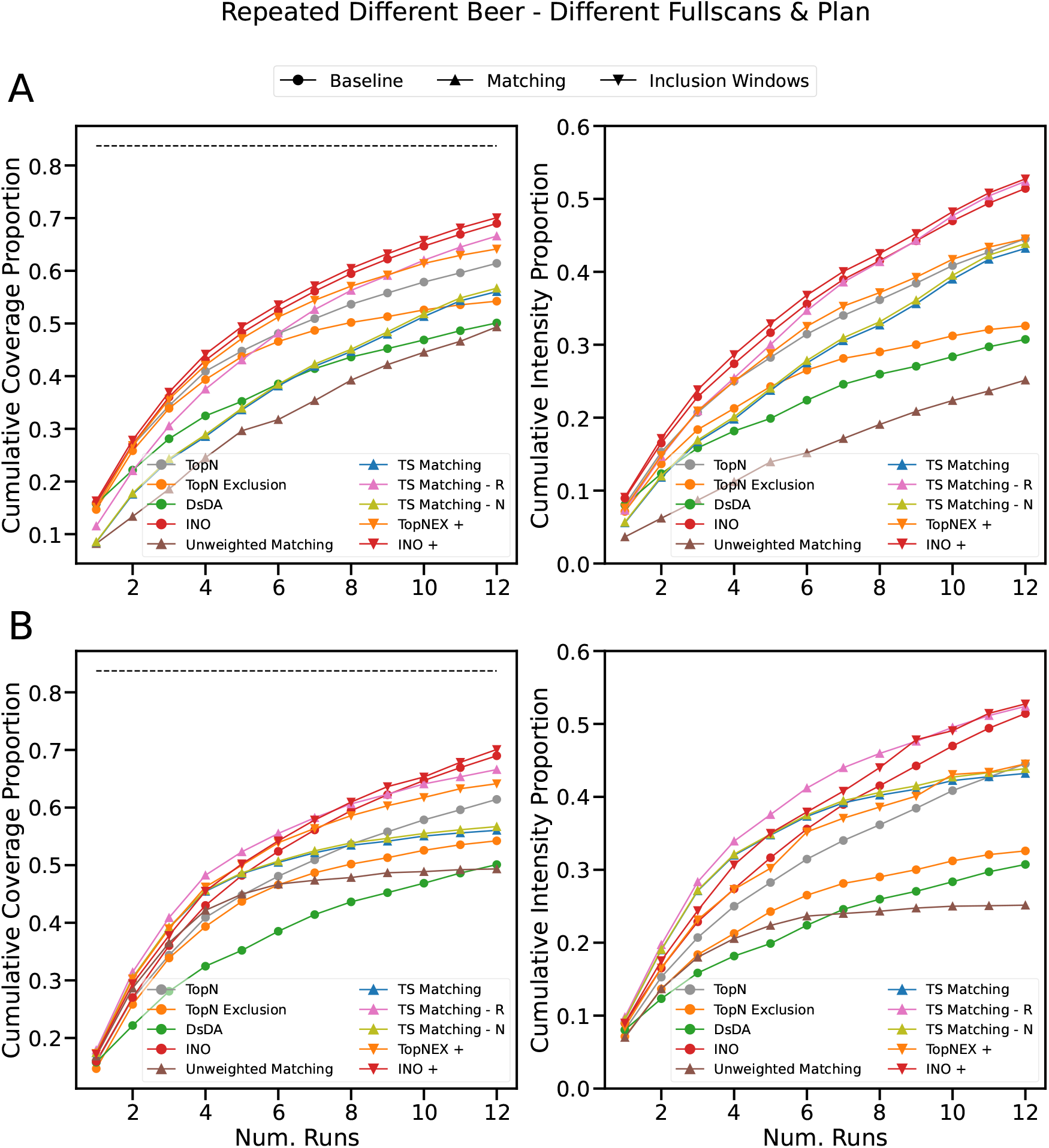
Experiment using four beers repeated three times each in round-robin order (1-2-3-4-1-2…). Two sets of twelve fullscans were used. One set was used to generate the target list for the matching algorithm, and the other was used for the simulations and the target list of the evaluation. XCMS was used to generate target lists for both the matching algorithm and the evaluation. The dotted line indicates the level of overlap between the target lists generated from the two sets of fullscans. **A:** shows performance over different runs in a single experiment. **B:** shows performance over separate experiments of different run numbers.

## 5 Discussion & Conclusion

In this work we have described a powerful maximum matching based technique for creating LC-MS/MS scan schedules for a given “target list” created from prior knowledge of the acquisition. We have used it both as a standalone “pre-scheduled” method and as an augment to existing Data-Dependent Acquisition methods by using it to generate inclusion windows. While the maximum matching problem has been mapped to the problem of LC-MS/MS acquisition control before [4], this is the first time it has been tested in terms of actually scheduling an acquisition, and we have also demonstrated the necessity of our improvements to the technique to make it practically applicable. We have also highlighted some of the trade-offs made between pure pre-scheduled methods and DDA methods.

Our experiments in Sections 4.1 and 4.2 showed impressive performance gains for the best fully pre-scheduled method, with it gaining over 20% on both coverage and intensity coverage in a simulated “best-case scenario” environment, theoretically enabling effectively completely comprehensive acquisitions. However, in Section 4.3, we saw that in a more realistic case its performance was similar to some of the other best-performing methods like Intensity Non-Overlap and TopN Exclusion augmented with inclusion windows generated by the matching workflow. This illustrates the main tradeoff made between pre-scheduled methods and DDA in that while pre-scheduled methods can create very sophisticated schedules which theoretically lead to highly comprehensive data acquisition, they may have difficulty realising this potential in practice.

In the same experiments we also demonstrated the usefulness of our improvements to the existing matching workflow. In all cases having access to more than one LC-MS/MS run allowed a more comprehensive data acquisition so the benefits of having a multi-run technique (which we introduced in Section 2.2) obviously follow. We also showed how multiple samples could be run as part of the same experiment. The “weighted matching” we introduced in Section 2.3 was justified both by the increased intensity coverage (the unweighted matching was often one of the worst performing methods at this metric) and by the fact it generally offered better coverage under the variations in the data we introduced in Section 4.3. Finally, the full assignments introduced in Section 2.4 was justified by the increase in robustness demonstrated in the same Section 4.3. The *recursive* variant of the matching workflow was the only method to show competitive performance with the other best methods.

For inclusion window generation, we decided to integrate this with our previous work on TopNEXt [16]. This was both because TopNEXt could be easily adapted to incorporate this functionality modularly and because it houses some of the current state-of-the-art in DDA acquisition methods (e.g. Intensity Non-Overlap, SmartRoI). While matching-generated inclusion windows did not seem to provide a significant benefit to Intensity Non-Overlap (which in general we found performed extremely well), they were a significant improvement on another established method, TopN Exclusion, often offering coverage increases of 10%. It should be noted that we generated our inclusion windows by assuming we knew the maximum number of runs in advance (as this is how our pre-scheduled method operates). In future, it may be better to generate inclusion windows greedily after every run to properly leverage the advantages of DDA.

However, this also illustrates one of the major trade-offs between DDA and pre-scheduled methods: DDA requires access to dynamic instrument control. If you cannot control the instrument in real-time then you cannot implement a DDA method by definition. Pre-scheduled methods, conversely, allow schedules to be translated beforehand into whichever format the instrument recognises. This may be one of the reasons why there is relatively little research on new DDA methods outside of frameworks which allow in-silico simulation like ViMMS [29]. While our matching-based method benefits from dynamic instrument control, because it allows it to run fully automatically and hybridisation with DDA requires dynamic control, at its core it is still a pre-scheduled method, so it can be used as-is. However, our experiments show the limitations of this totally offline approach, and we predict as this field develops dynamic algorithms will become more important. Our matching workflow may also integrate with this trend, for example by the use of a dynamic algorithm [2], which would make it more robust and allow it better performance in practice.

Another one of the strengths of our matching workflow is that it can be composed with a completely arbitrary target list of the user’s choice. The matching algorithm merely resolves an optimal schedule across this list of targets. We used peak-picking to generate the target list, but it is equally possible to, for example, integrate the matching algorithm’s scheduling with a CAMERA-style acquisition workflow [14, 32]. Alternatively, a hypothetical future piece of software could assign a “utility weight” to each edge, expressing the ultimate usefulness of that acquisition as a single numerical estimate. It would then be possible to plan a number of runs to obtain, say, 90% coverage or utility coverage in a completely closed-loop way. More complex problem formulations like stable marriage [7] or a constraint programming formulation [27] may also allow encoding more complex trade-offs between utility values.

The modularity of our method is also important because we have seen that the effectiveness of our workflow depends heavily on the peak-picking. The differences between the experiments in Sections 4.1, and 4.3 are essentially a product of different peak-picking assumptions in the evaluation, and as we have seen, the differences are quite large. While conclusions from our evaluation can be supplemented by metabolite annotations from experimental data (and we hope to see such validations in future) this is also potentially a core bottleneck of the acquisition workflow we have presented (in terms of generating spurious targets, improperly aligning data, etc). The modularity of our approach means, however, that future improvements in peak-picking software can be easily integrated into the workflow.

Overall, we have shown a promising new technique for LC-MS/MS acquisition software control which competes with many state-of-the-art methods. This maximum matching based technique has many promising directions for future development, but can also be deployed as-is either using dynamic instrument control or as a completely offline workflow.

## Authors’ contributions

RM performed the bulk of design and implementation of the methods, conducted computational experiments and wrote the initial draft of the manuscript. SW performed all labwork in seed data collection. JW was involved in implementation improvements to ViMMS to facilitate this project and gave continuous feedback on the project. VD, RD and KB supervised the project. Additionally, VD contributed substantially to details of method design and RD contributed substantially to experiment design and to obtaining time on the relevant lab instrument. All authors approved the final manuscript.

## Availability of data and materials

The matching fragmentation strategies are implemented as part of the ViMMS framework and the latest version can be found at https://github.com/glasgowcompbio/vimms. A stable version used to produce our results can be found at 10.5281/zenodo.6394524. Data can be found at 10.5525/gla.researchdata.1877.

## Declarations

### Ethics approval and consent to participate

Not applicable.

### Consent for publication

Not applicable.

