## Supplementary Information for "Comprehensive LC-MS/MS Data Acquisition in Metabolomics via Maximum Bipartite Matching"

#### 1 Beer Samples

In our experiments we used four different beers that were originally collected in [1]. The specific beers used have been reproduced from [1] in Supplementary Table 1. For the repeated single beer experiment we used sample 1 — all four were used for the different beers experiment.

| Index | Name | Type |
|---|---|---|
| 1 | Raspberry Sour by Vault City | Sour |
| 2 | Cacao & Hazelnut Broken Dream Twisted Breakfast Stout by Siren | Stout |
| 3 | Tennents | Lager |
| 4 | Life and Death by Vocation | IPA |

Table 1: Names and types of beer used to seed experiments.

#### 2 Resynchronisation

Pre-scheduled methods require a known schedule of scan times and levels ahead of time. However, in practice an LC-MS/MS setup may produce scans at unexpected times — for example there may be variation in how long it takes to collect a sufficient number of ions for a scan. In the most extreme case the schedule could become completely desynchronised.

To run our pre-scheduled matching on a lab instrument in future we would require some means to resynchronise the planned schedule with the actual scan times observed. We have therefore implemented dynamic re-scheduling of a scan schedule for our pre-scheduled controllers. Additionally, this functionality is implemented with a generic pre-scheduled controller so it can be used interchangeably with the pre-scheduled matching, DsDA, or another subclass.

The implementation of the resynchonisation holds the schedule of scans in a buffer and adds and removes scans to keep times synchronised. A "filler" MS1 or MS2 scan is added if the next scan's planned RT exceeds the current RT by more than a user-specified parameter — the scan type is chosen to be the longest scan that will fit. "Filler" MS2 scans target the same target as the nearest MS2 in the schedule. Conversely, we cancel the next scan if we are too close to the intended RT of its successor — again, a user-specified

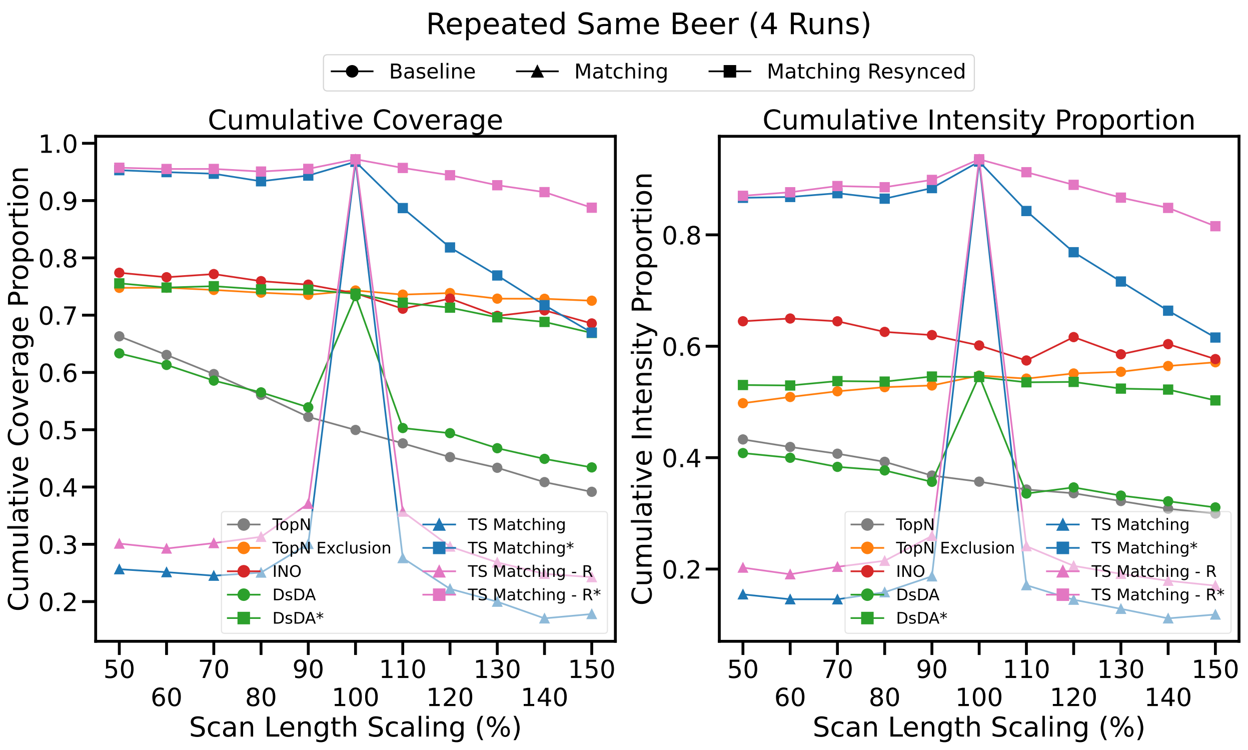


Figure 1: Coverage and intensity coverage for experiments of the same beer being run consecutively for four iterations and with XCMS peak-picking. Each point on the x-axis represents a separate experiment where scan lengths were scaled to x%. Squares indicate a resynchronised version of the matching method which can account for the unexpected scan lengths.

parameter defines how close is too close. If neither of these conditions are met, then we remove the next scan in the buffer and queue it on the instrument.

As a test of this functionality we conducted an experiment where scan times were deliberately completely desynchronised from the times pre-scheduled methods expected. Each x-axis value in Supplementary Figure 1 represents a same beer repeated experiment of four iterations where scan lengths were multiplied by a different constant factor, ensuring that the schedule quickly becomes completely desynchronised as some displacement in the same direction is added per each scan queued.

In general, method performance trends downwards as the scan length increases, decreasing the number of scans available. However, for the baseline pre-scheduled methods, this trend is dominated by the effect of the schedule becoming desynchronised. For the matching methods, performance drops massively as scan times become desynchronised in either direction. Compared to the matching methods, DsDA has lower performance with a synchronised schedule, but retains a reasonable level of performance when desynchronised in either direction, perhaps suggesting that it is "self-healing" because it continues to use results of runs for planning as they arrive. Conversely, the resynchronised matching methods do not have this performance drop-off and remain largely constant other than losing some performance to having fewer scans available as scan lengths shorten.

There are some caveats with interpreting these experiments. Pre-scheduled methods cease to send scans when they have executed their schedule, causing the acquisition to end earlier with shorter scan lengths, and longer scan lengths may cause some peaks to not be detectable in the simulation. Because the evaluation only counts those peaks which were detectable in the fragmentation runs, these factors could cause some values to be artificially higher.

Additionally, there is a slight drop-off in resynchronised matching strategy performance with shorter scan times, which should theoretically allow better performance. This may suggest the resynchronisation parameter choices do not allow the schedule to be totally mapped to the new scan times. Still, it is a minor difference and a real-world scenario should have less systematic scan-time biases, which should minimise the impact of parameter choice.

Despite these caveats we can conclude that this resynchronisation procedure is effective in correcting the systematic scan time biases we tested, and it should be suitable for less severe conditions in reality.

### 3 DsDA Implementation

To interface ViMMS with DsDA for the purposes of our comparison we chose to use the Python *subprocess* module to invoke a version of the DsDA script modified to take command-line parameters. A complete Python re-implementation of DsDA would risk errors in translation and thus a spurious comparison. The Python-based ViMMS DsDA controller and the DsDA R script must take turns running, so on invocation the parent Python process awaits the completion of the R process. Previous work [2] initialised DsDA manually and had communication performed by ad-hoc polling of output directories, but this cannot be scaled for future comparisons (e.g. suppose a future comparison must run 10 separate methods individually, or that unrelated files are being written to the output directory).

With our new implementation, communication between ViMMS and DsDA is still performed via the file system — ViMMS writes an .mzML and DsDA is informed, then DsDA writes a scan schedule for the next LC-MS/MS run and ViMMS is informed. However, the update happens by a socket connection passing the filename (a reasonable alternative would be to have them poll a shared temporary file). A Python *DsDAProcess* object wraps this connection and allows DsDA to be used like any other ViMMS controller. Note that using the original DsDA implementation in this way also motivated our evaluation to use XCMS — because DsDA uses XCMS internally, this was necessary to ensure a fair comparison.

### 4 MZMine Parameters and Experiments

The results in main paper Section 4 showed XCMS results with the parameters in Supplementary Table 2, which have a low quality-filter. We used XCMS both because it was fairer to DsDA (which uses XCMS internally) and the low quality-filter made the peak scheduling problem harder and allowed us to show that the matching algorithm is in theory capable of resolving a schedule for a very high number of peaks at once.

| **Centwave Parameters** | | **Join Aligner Parameters** | |
|---|---|---|---|
| ppm | 15 | mzVsRtBalance | 10 |
| peakwidth | (15, 80) | absMz | 0.2 |
| snthresh | 5 | absRt | 15 |
| noise | 1000 | kNN | 10 |
| prefilter | (3, 500) | | |
| mzdiff | 0.001 | | |

Table 2: Parameters used for XCMS peak-picking and alignment.

However, it is also interesting to ask what results are like for the restrictive MZMine parameter set from the TopNEXt publication [1], which produces a much smaller but higher-quality set of peaks. To illustrate this, Table 1 from the main paper shows us that there is a roughly 3:1 ratio between XCMS peaks and MZMine (Restrictive) peaks. This is a bit lower than the ratio between the two MZMine parameter sets on the TopNEXt experiments' dataset because the number of peaks in the more restrictive parameter set drops off less sharply when given the negative ionisation mode data we have used this time. Nonetheless, results from the two different peak-picking parameter sets are very different and produce very different results when evaluating a fragmentation strategy. The restrictive parameter set has been reproduced from [1] (Supplementary Section 2) in Supplementary Table 3

Supplementary Figure 2 shows the results of this procedure for the same beer, single fullscan case (i.e. equivalent to main paper Figure 6 but using different peak-picking to construct target lists). It is immediately apparent, like with the comparison between restrictive and permissive parameters from the work on TopNEXt [1], that the scale on these plots is very different to those generated with our XCMS peak-picking setup.

| MZMine Parameters (Restrictive) | |
|---|---|
| **Raw Data Import** | |
| **Crop Filter** | |
| Retention time | 0.5 to 30 minutes |
| $m/z$ | 50 to 1060 Da |
| **Mass Detection (MS1)** | |
| Mass detector | Centroid |
| Noise level | 1000 |
| **Mass Detection (MS2)** | |
| Mass detector | Centroid |
| Noise level | 0 |
| **ADAP Chromatogram Builder** | |
| MS level | 1 |
| Min group size in # of scans | 3 |
| Group intensity threshold | *2000* |
| Min highest intensity | 5000 |
| $m/z$ tolerance$^{\dagger}$ | $10^{(-8)}$ Da or 10.0 ppm |
| **Chromatogram Deconvolution** | |
| Algorithm | Wavelets (ADAP) |
| $S/N$ threshold | *3* |
| $S/N$ estimator | Intensity window SN |
| Min feature height | 5000 |
| Coefficient/area threshold | 1 |
| Peak duration range | *1.0 to 7.00* |
| rt wavelet range | *1.00 to 5.00* |
| $m/z$ centre calculation | Median |
| Range for MS2 scan pairing ($m/z$, RT) | 0.01 Da, 0.5 minutes |
| **Isotopic Peaks Grouper** | |
| $m/z$ tolerance$^{\dagger}$ | $10^{(-8)}$ Da or 5.0 ppm |
| Retention time tolerance | 0.1, absolute (minutes) |
| Monotonic shape | False |
| Maximum charge | 2 |
| Representative isotope | *Lowest m/z* |
| **Join Aligner** | |
| $m/z$ tolerance$^{\dagger}$ | $10^{(-8)}$ Da or 10.0 ppm |
| Weight for $m/z$ | 80 |
| Retention time tolerance | *0.5, absolute (minutes)* |
| Weight for RT | 20 |
| Require same charge state | False |
| Compare isotope pattern | False |
| Compare spectra similarity | False |
| **Export to CSV** | |

Table 3: Our set of "restrictive" MZMine 2 batch mode parameters, reproduced from [1].

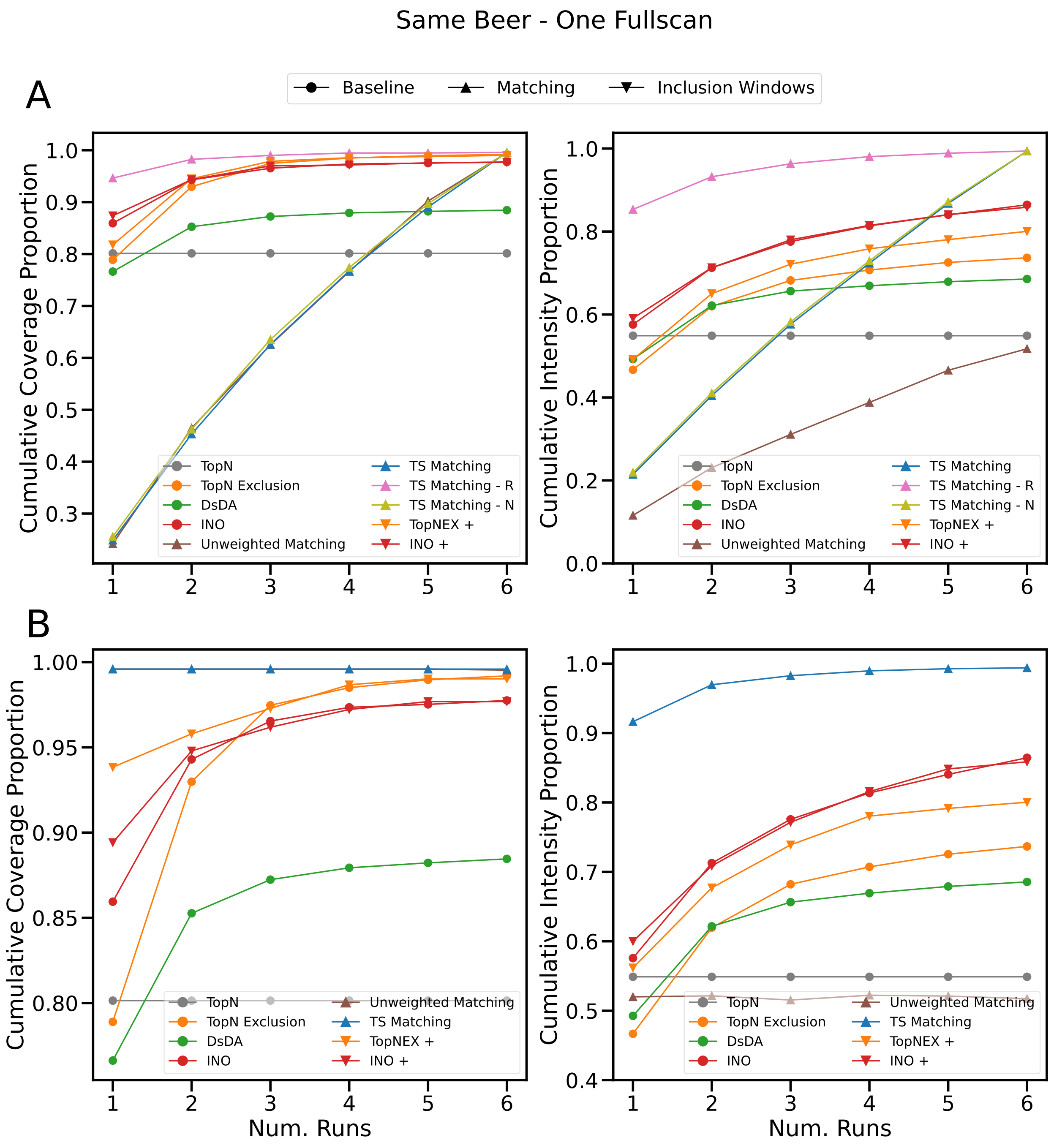


Figure 2: Experiment using the same beer repeated for six samples. A single fullscan was used to generate simulations and target lists, and MZMine (restrictive parameter set) was used to generate target lists for both the matching algorithm and the evaluation. **A:** shows performance over different runs in a single experiment. **B:** shows performance over separate experiments of different run numbers.

Essentially every method (minus TopN and DsDA) gets to 100% coverage and over 70% intensity coverage. The two-step matching is able to get to roughly 100% coverage and over 90% coverage in only a single run.

Broadly, we see some of the trends we have come to expect from this data. Pre-scheduled matching is very good when it has perfect knowledge of what is coming. Inclusion windows improve both Intensity Non-

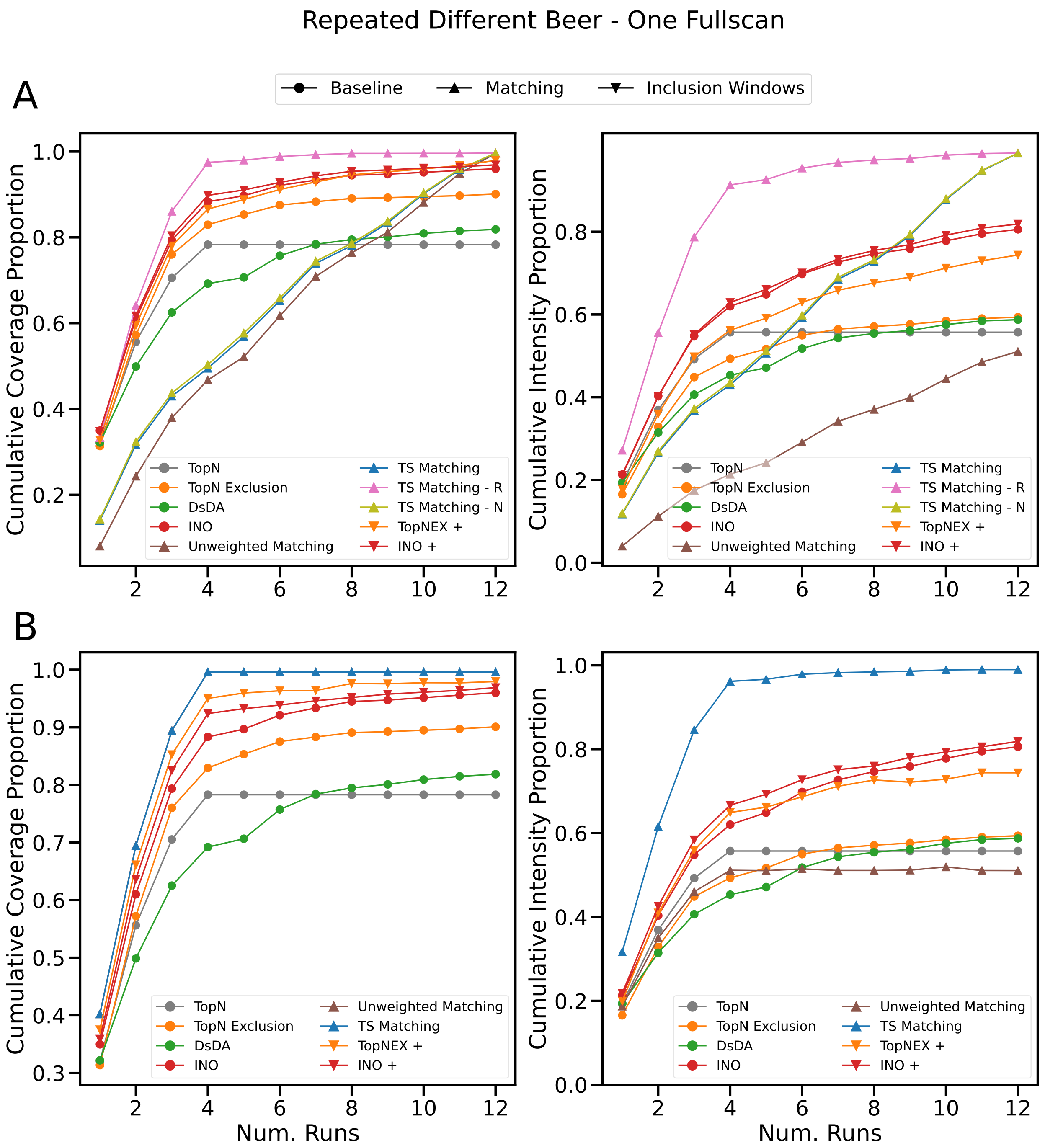


Figure 3: Experiment using four beers repeated three times each in round-robin order (1-2-3-4-1-2...). A single fullscan was used to generate simulations and target lists, and MZMine (restrictive parameter set) was used to generate target lists for both the matching algorithm and the evaluation. **A:** shows performance over different runs in a single experiment. **B:** shows performance over separate experiments of different run numbers.

Overlap and TopN Exclusion, but moreso for TopN Exclusion. And TopN has very poor performance when there is no variation in the underlying data (although it is worth remembering there will always be variation in reality). Compared to the XCMS-based experiments, both variants of Intensity Non-Overlap show somewhat poorer performance in coverage compared to TopN Exclusion with inclusion windows, but Intensity Non-Overlap maintains its intensity coverage advantage. One notable change is that the performance of the unweighted matching looks worse relative to Figure 6: but this is a function of the fact that it is generally easier for other methods to obtain coverage.

At a glance the most interesting trend is that DsDA appears to saturate below 90% coverage and 70% intensity coverage. This could be because its scoring function is prone to saturation or because of the mismatch between the XCMS and MZMine parameter settings (we did not adjust between these experiments) or some combination of both. It is therefore important not to read too deeply into DsDA's performance in this section, but it is notable that it (albeit like every other method) manages to target the majority of the evaluation target list in a scenario where it has a different target list due to using XCMS. These difficulties with comparisons using DsDA in this scenario also speak to the need for a more integrated platform for testing and running LC-MS/MS fragmentation strategies (i.e. continued development of ViMMS).

Examining Supplementary Figure 3 we also observe the same basic trends but for the 4-3 repeated different beers experiment (with re-used fullscans). The pre-scheduled matching gets effectively total coverage and almost total intensity coverage (in the case of the two-step matching) once it has seen all four samples. Many methods again eventually converge to approx. 100% coverage, but likely due to the increased complexity of this experiment compared to the same beer case, the intensity coverage gap between the two-step matching and the next-best method has widened as well. We also see again that TopN Exclusion with inclusion windows is the next most effective method in coverage, but Intensity Non-Overlap outperforms it in intensity coverage. DsDA also seems to saturate after slightly outperforming TopN — both of these trends are similar to Figure 7.

Supplementary Figures 4 and 5 show the different fullscan scenario (using a different fullscan per run but sharing them between the matching algorithm and simulator) for the same beer and repeated different beer (4-3) cases respectively. These essentially reiterate the same basic trends we have seen in the main paper. Like main paper Figures 8 and 9 performance is more slowly gained over the total length of the experiment (due to more target peaks appearing with each run) but like the previous experiments using MZMine we also see that the numbers overall are higher. The pre-scheduled matching still has very strong performance given that it knows what is coming, but the inclusion window methods and base Intensity Non-Overlap track it more closely due to the overall number of peaks being lower. We also see the performance of DsDA and TopN Exclusion falling relative to TopN in Supplementary Figure 5 as we saw for the repeated different beer experiment before.

Finally, we will consider the case where two seperate sets are used to create the target list for the matching algorithm and for the simulator and evaluation, but under the restrictive MZMine parameter set. Supplementary Figure 6 shows this for the same beer case. Like main paper Figure 10, the performance of the matching-based methods drops severely in this more realistic scenario where they do not know exactly what is coming. However, while using the XCMS parameters the recursive pre-scheduled matching was a reasonable competitor for best method, here it is behind both variants of TopN Exclusion and both variants of Intensity Non-Overlap in coverage.

Additionally, the other versions of the pre-scheduled matching perform only around as well as DsDA, well behind TopN (DsDA is also doing more poorly because of the smaller number of peaks). This is because without a large number of peaks to resolve a schedule for (and with only a relatively small number of higher-quality peaks in the target list) the advantages of the pre-scheduled matching are less prominent. 80% is a relatively high coverage score, but Intensity Non-Overlap reaches higher, at 90%. The matching method fares somewhat better in intensity coverage (beating base TopN Exclusion and competing with the inclusion window variant) but Intensity Non-Overlap is still the best performer.

This drop in matching-based performance also applies to the inclusion window methods. The two Intensity Non-Overlap variants are essentially identical in performance, and while adding inclusion windows is still a significant boost to TopN Exclusion, it no longer allows it to outperform Intensity Non-Overlap in coverage. One interesting thing to note, however, is that the recursive variant of the pre-scheduled two-step matching is above the dotted line, even though this means it is hitting things not included in the target list. This may suggest the width of the isolation window is large enough that it hits targets that MZMine has not managed to align together (possibly in dense areas of the data).

Supplementary Figure 7 also reiterates a lot of the same basic patterns. Intensity Non-Overlap (either

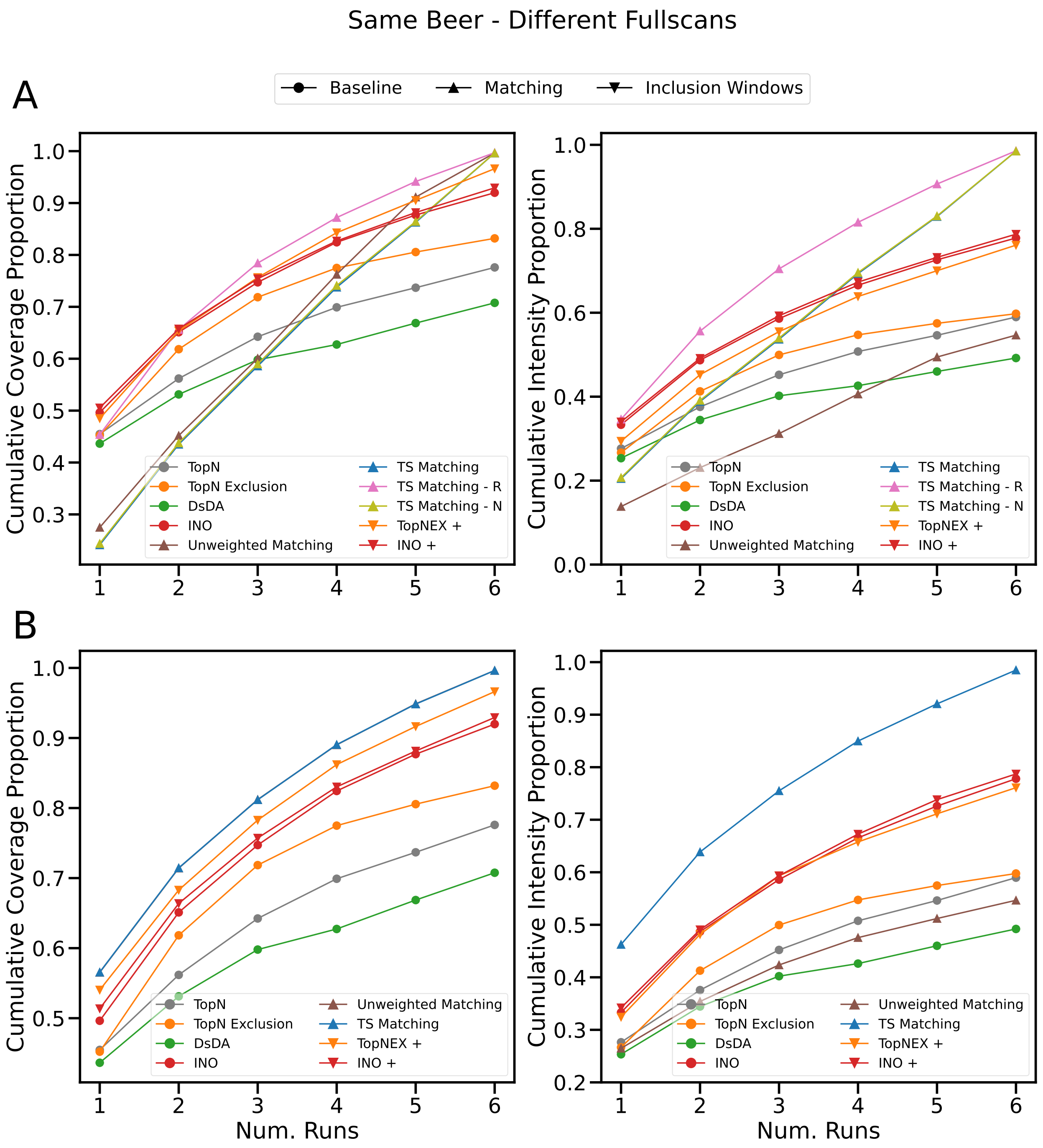


Figure 4: Experiment using the same beer repeated for six samples. A different fullscan for each of the six runs was used to generate simulations and target lists, and MZMine (restrictive parameter set) was used to generate target lists for both the matching algorithm and the evaluation. **A:** shows performance over different runs in a single experiment. **B:** shows performance over separate experiments of different run numbers.

variant) remains the most effective method, pre-scheduled methods suffer relative to other methods because of the smaller peak number (though the recursive variant is still effective) and the gain for inclusion windows

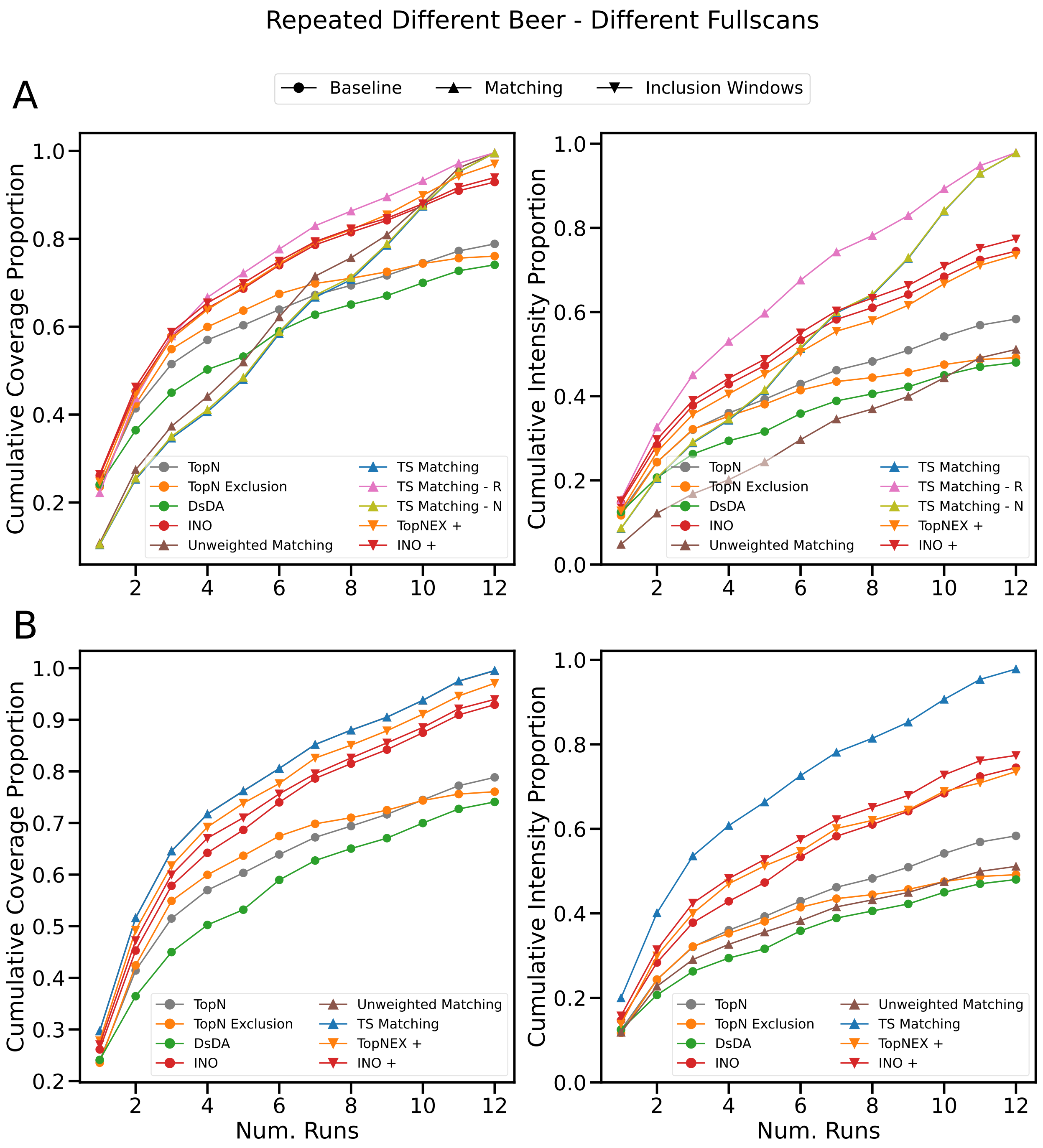


Figure 5: Experiment using four beers repeated three times each in round-robin order (1-2-3-4-1-2...). A different fullscan for each of the twelve runs was used to generate simulations and target lists, and MZMine (restrictive parameter set) was used to generate target lists for both the matching algorithm and the evaluation. **A:** shows performance over different runs in a single experiment. **B:** shows performance over separate experiments of different run numbers.

is also smaller relative to when the data was perfectly known in advance (though TopN Exclusion still benefits substantially). One notable change from Supplementary Figure 6 is that the recursive two-step matching

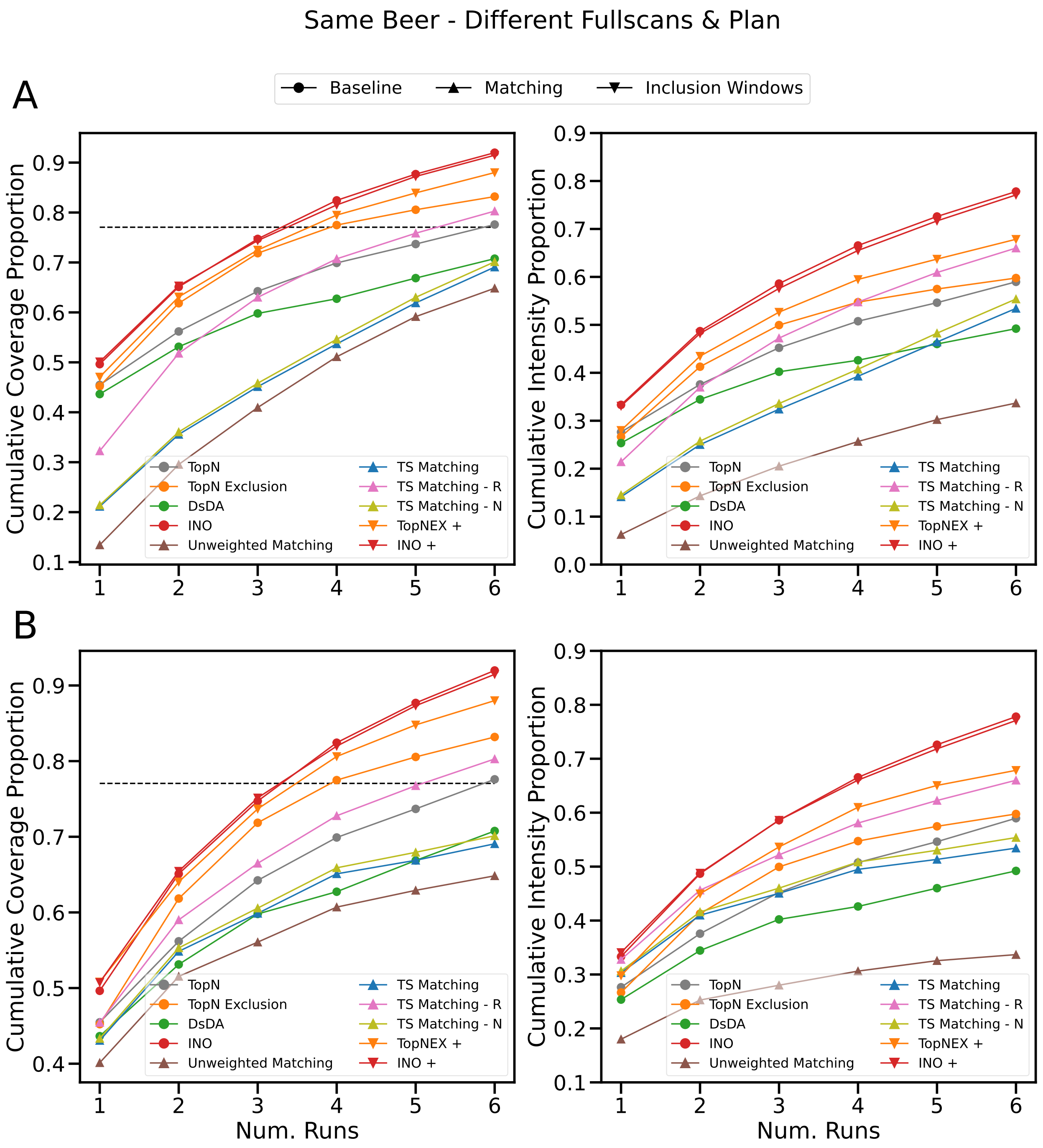


Figure 6: Experiment using the same beer repeated for six samples. Two sets of six fullscans were used. One set was used to generate the target list for the matching algorithm, and the other was used for the simulations and the target list of the evaluation. MZMine (restrictive parameter set) was used to generate target lists for both the matching algorithm and the evaluation. The dotted line indicates the level of overlap between the target lists generated from the two sets of fullscans. **A:** shows performance over different runs in a single experiment. **B:** shows performance over separate experiments of different run numbers.

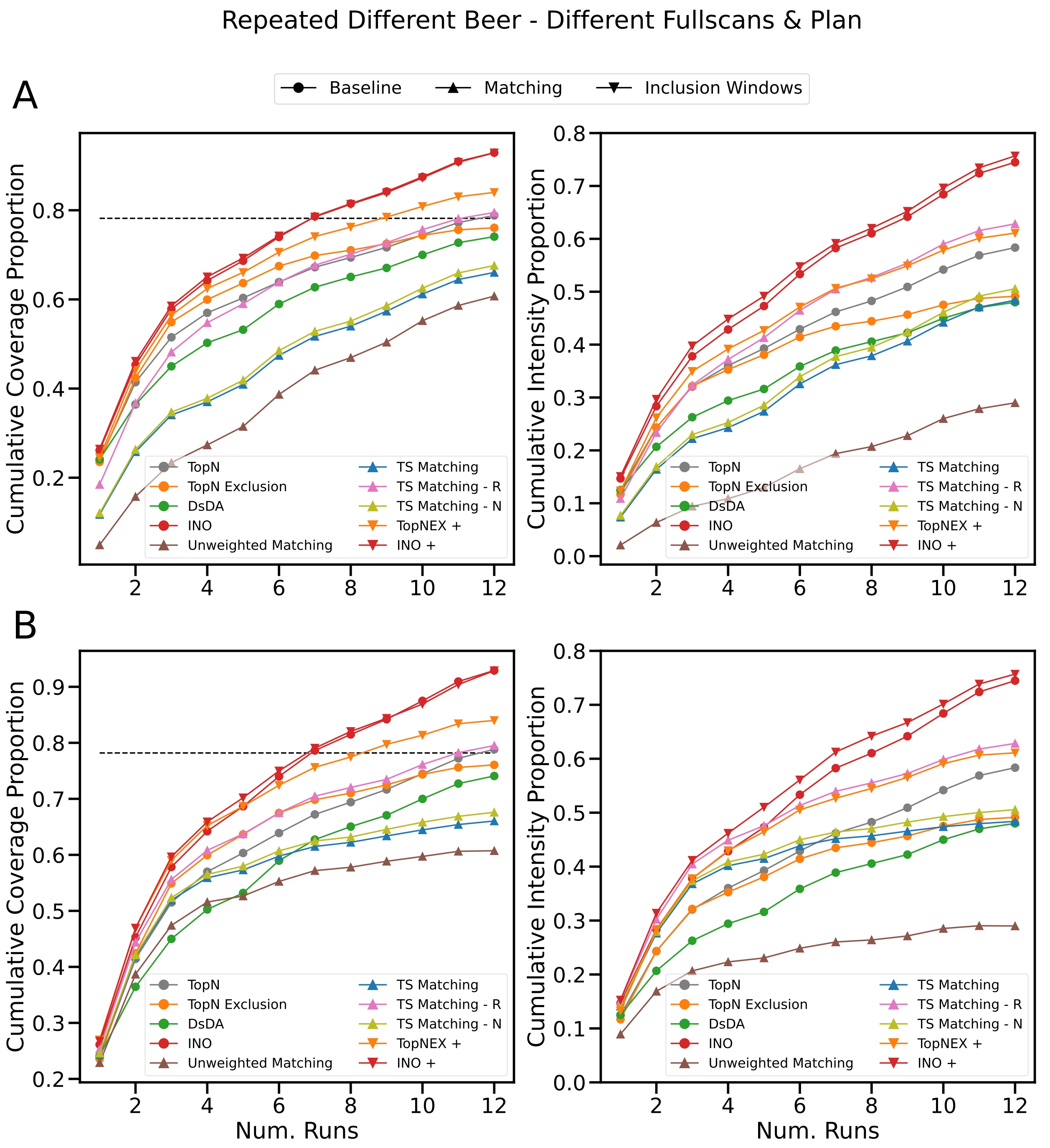


Figure 7: Experiment using four beers repeated three times each in round-robin order (1-2-3-4-1-2...). Two sets of twelve fullscans were used. One set was used to generate the target list for the matching algorithm, and the other was used for the simulations and the target list of the evaluation. MZMine (restrictive parameter set) was used to generate target lists for both the matching algorithm and the evaluation. The dotted line indicates the level of overlap between the target lists generated from the two sets of fullscans. **A:** shows performance over different runs in a single experiment. **B:** shows performance over separate experiments of different run numbers.

variant outperforms base TopN Exclusion in coverage (as we have seen, TopN Exclusion particularly struggles with this experiment design using multiple sample types). DsDA is also closer to reaching base TopN in performance, likely thanks to the increased number of runs.

Overall, we have seen that the lower number of peaks produced by the MZMine restrictive parameter-set has strong effects on the best choice of fragmentation strategy. While it moves the entire scale of the results up so that in theory only one or two runs per sample are enough for a complete acquisition, the pre-scheduled matching struggles in an environment where reality differs from its target list. While its performance remains competitive, this underscores the tradeoff being made between DDA and pre-scheduled methods. Additionally, we have confirmed our earlier results from our work on TopNEXt [1] by observing that Intensity Non-Overlap is a very effective method (in terms of both coverage and intensity coverage) at lower peak numbers. Unfortunately, it does not seem that (unlike TopN Exclusion) matching-generated inclusion windows improve it much, if at all, thanks to its more complex behaviour — although it will require more testing to see whether this was due to the base behaviour of Intensity Non-Overlap or due to its combination with SmartRoI for these experiments.

### 5 Timings

One of the factors that might affect the performance of a pre-scheduled method is how much time elapses between the collection of its representative data and the actual acquisition that has been planned. While ideally runs would be conducted directly one after another, processing overhead may increase the gap between these two steps. For this reason, we timed each part of the matching workflow while generating data for our experiments, and present the timings in Supplementary Table 4. These timings were collected on an ordinary desktop PC with an Intel(R) Core(TM) i7-10700 2.90 GHz CPU and 32GB of RAM. Importantly, these timings were not collected in a controlled environment (i.e. other programs were running on the machine and some parts of the matching procedure which we were collecting separate timings for ran in parallel) nor were they averaged across replicates. Consequently, the timings should *only* be interpreted as a rough guide as to how much time the workflow will need, not as precise algorithmic timings.

In general there is no significant difference between XCMS and MZMine (with restrictive parameters) when it comes to processing time for the workflow, despite the very different number of peaks. The longest-running part of the workflow is the "Scan Creation" step, where scans from the representative data are interpolated to give expected intensity values for each scan in the schedule. In the majority of cases it is slower than all other steps combined. Scan creation is also notably slower when being used in the cases where we had a different fullscan per run because it simply had more files to process.

The scan creation subroutine uses ViMMS' implementation of RoI-building, is written in pure Python and is slow as a result of the very large amount of data in .mzMLs there is to process. However, the implementation of this step could be improved in future (for example, XCMS' implementation of RoI-building calls linked C libraries despite the software primarily being written in R). With that being said, this step is actually significantly less of a time bottleneck in our workflow than peak-picking — for example, MZMine had to be left to run over the course of several days to generate all of the peak-picked files we used.

While the two-step matching generally has a runtime twice or thrice as high as the unweighted matching, none of these runtimes exceed 3 minutes. To add context to this number, the scan creation times often exceed ten times this amount and it is not unusual for the length of a single LC-MS/MS metabolomics run to be half an hour. Given that we have seen that the unweighted matching is significantly less effective than the two-step matching, this seems like a small price to pay. Still, if this was too slow, it is possible to apply heuristics like only keeping the $n$ edges with the highest weight for each vertex (potentially at the expense of results quality).

The full assignment step does, on the other hand, consume a significant amount of processing time when using *recursive* mode, occasionally exceeding the length of scan creation time and reaching nearly half an hour in one case. While the processing time for *nearest* is negligible, we also saw in our experiments that it barely registered an improvement in coverage and intensity coverage. It may be desirable in future to have a full assignment heuristic which redundantly spreads targets across runs like *recursive* but which is less processing-intensive.

| Same Beer | | | | |
|---|---|---|---|---|
| | Re-used Fullscans | | Unique Fullscans | |
| | XCMS | MZMine | XCMS | MZMine |
| Scan Creation | 6.5 mins | 6.3 mins | 32 mins | 31.8 mins |
| Unweighted Matching | 17 secs | 30 secs | 21 secs | 29 secs |
| Two-Step Matching | 49 secs | 1.3 mins | 56 secs | 1.3 mins |
| Recursive Assignment | 5.4 mins | 13.8 mins | 4.9 mins | 8.7 mins |

| Repeated Different Beer | | | | |
|---|---|---|---|---|
| | Re-used Fullscans | | Unique Fullscans | |
| | XCMS | MZMine | XCMS | MZMine |
| Scan Creation | 24.8 mins | 24.7 mins | 69.1 mins | 68 mins |
| Unweighted Matching | 48 secs | 1.1 mins | 45 secs | 1.1 mins |
| Two-Step Matching | 2.1 mins | 3 mins | 2.5 mins | 2.9 mins |
| Recursive Assignment | 13.8 mins | 26.2 mins | 11.8 mins | 20.1 mins |

Table 4: Times elapsed for different stages of the matching process during the experiments in Section 4 and Supplementary Section 4 (some stages with negligible runtime have been omitted). Scan creation was run separately for the unweighted and two-step matching so times for those two cases are averaged.
